# Deciphering and modelling the TGF-β signalling interplays specifying the dorsal-ventral axis of the sea urchin embryo

**DOI:** 10.1101/2020.02.26.966556

**Authors:** Swann Floc’hlay, Maria Dolores Molina, Céline Hernandez, Emmanuel Haillot, Morgane Thomas-Chollier, Thierry Lepage, Denis Thieffry

**Affiliations:** Institut de Biologie de l’ENS (IBENS), École Normale Supérieure, CNRS, INSERM, Université PSL, 75005 Paris, France.; Institut Biologie Valrose, Université Côte d’Azur, Nice, France.; Institut Universitaire de France (IUF), 75005 Paris, France.

**Keywords:** *Paracentrotus lividus*, embryo development, logical model, gene regulatory network, Nodal pathway, BMP pathway.

## Abstract

During sea urchin development, secretion of Nodal and BMP2/4 ligands and their antagonists Lefty and Chordin from a ventral organizer region specifies the ventral and dorsal territories. This process relies on a complex interplay between the Nodal and BMP pathways through numerous regulatory circuits. To decipher the interplay between these pathways, we used a combination of treatments with recombinant Nodal and BMP2/4 proteins and a computational modelling approach. We assembled a logical model focusing on cell responses to signalling inputs along the dorsal-ventral axis, which was extended to cover ligand diffusion and enable multicellular simulations. Our model simulations accurately recapitulate gene expression in wild type embryos, accounting for the specification of ventral ectoderm, ciliary band and dorsal ectoderm. Our model simulations further recapitulate various morphant phenotypes, reveals a dominance of the BMP pathway over the Nodal pathway, and stresses the crucial impact of the rate of Smad activation in D/V patterning. These results emphasise the key role of the mutual antagonism between the Nodal and BMP2/4 pathways in driving early dorsal-ventral patterning of the sea urchin embryo.

**Summary Statement:** We propose a predictive computational model of the regulatory network controlling the dorsal-ventral axis specification in sea urchin embryos, and highlight key features of Nodal and BMP antagonism.

## Introduction

During embryonic development, cell fate is specified by transcription factors activated in response to instructive signals. Regulatory interactions between signalling molecules and their target genes form networks, called Gene Regulatory Networks (GRN) (Arnone and Davidson, 1997). Deciphering such GRNs is a key for developmental biologists to understand how information encoded in the genome is translated into cell fates, then into tissues and organs, and how morphological form and body plan can emerge from the linear DNA sequence of the chromosomes (Levine and Davidson, 2005). Noteworthy, the gene regulatory network orchestrating the morphogenesis of the ectoderm along the dorsal-ventral axis of the embryo of the model sea urchin *Paracentrotus lividus* has started to be uncovered in great detail (Haillot et al., 2015; Lapraz et al., 2015; Li et al., 2012; Li et al., 2013; Li et al., 2014; Molina et al., 2018; Range et al., 2007; Saudemont et al., 2010; Su et al., 2009 and reviewed in Molina et al., 2013).

The ectoderm of the sea urchin larva is constituted of two opposite ventral and dorsal territories, separated by a central ciliary band (Fig. 1A):

**Figure 1.**
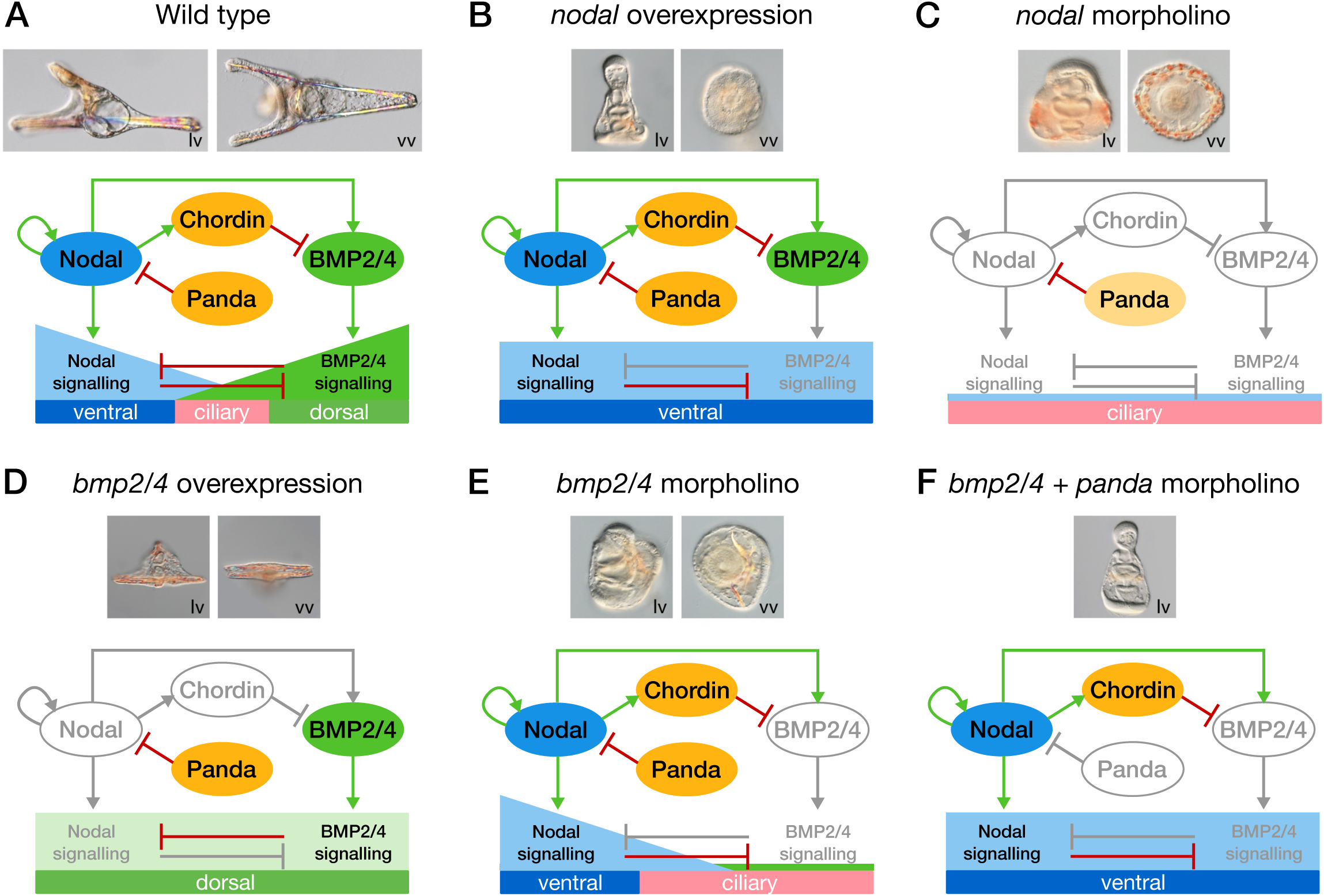
Panda, Nodal and BMP signalling direct patterning of the Dorsal/Ventral axis of the sea urchin embryo. Summary of the morphological phenotypes and identity of the ectodermal territories of wild- type (control) embryos and embryos following perturbations of Nodal or BMP signalling. (A) In control embryos, the balance between Nodal and BMP signalling patterns the ectoderm in three main territories: Nodal signalling specifies the ventral ectoderm, BMP signalling specifies the dorsal ectoderm, while a ciliary band develops at the interface between them. (B) The whole ectoderm differentiates into ventral territory when *nodal* is overexpressed. (C) Both Nodal and BMP24 signalling are absent in Nodal morphants, which give rise to an expanded large ciliary band. (D) Following BMP2/4 overexpression, all the ectoderm acquires dorsal identity. (E) In contrast, after BMP2/4 inhibition, ventral territories are not perturbed but an ectopic ciliary band develop in place of the presumptive dorsal ectoderm. (F) Simultaneous perturbation of both the TGF-β Panda and BMP2/4 signalling allows the expansion of Nodal signalling to the whole territory and the ventralisation of the ectoderm. The genes, proteins or interactions that are inactive following each perturbation are denoted in light grey. Activation and inhibition interactions are respectively shown by green and red arrows. lv, lateral view. vv, vegetal view.

The ventral ectoderm is the territory at the centre of which the mouth will be formed. Specification of the ventral ectoderm critically relies on signalling by Nodal, a secreted growth factor of the TGF-β family (Duboc et al., 2004; reviewed in Molina et al., 2013). The expression of *nodal* is turned on by maternal factors, while Nodal stimulates and maintains its own expression through a positive feedback circuit (Bolouri and Davidson 2010; Range et al., 2007; Range and Lepage, 2011). Nodal is zygotically expressed and is thought to dimerise with another TGF-β ligand maternally expressed called Univin (Range et al., 2007); the Nodal-Univin heterodimer promotes Alk4/5/7 signalling and the activation of Smad2/3 together with Smad4. The ventral ectoderm boundary is thought to be positioned by the activity of the product of the Nodal target gene *lefty*, which prevents the expansion of *nodal* expression beyond the ventral ectoderm region via a diffusion-repression mechanism (Chen et al., 2004; Cheng et al., 2004; Duboc et al., 2008; Molina et al., 2013; Sakuma et al., 2002).

The ciliary band ectoderm is a proneural territory located between the ventral and dorsal ectoderm (Angerer et al., 2011). The ciliary band is made of prototypical cuboidal epithelial cells and runs along the arms of the pluteus larva. Unlike the specification of the ventral and the dorsal ectoderm, which actively requires TGF-β signalling, the specification of the ciliary band tissue does not rely on Nodal or BMP signalling, and this tissue develops as a “default” state of the ectoderm in the absence of these signals (Saudemont et al., 2010).

The dorsal ectoderm is the territory that will differentiate into the apex of the pluteus larva. Its specification relies on the diffusion of ventrally synthesized BMP2/4, which promotes dorsal fates by activating phosphorylation of Smad1/5/8 via the activation of the BMP type I receptors Alk1/2 and Alk3/6. The inhibition of BMP signalling, on the ventral side, and the translocation of BMP2/4, to the dorsal side, require the product of the *chordin* gene, which is activated in the ventral ectoderm downstream of Nodal signalling (Lapraz et al., 2009). Glypican5 is expressed downstream of BMP2/4 signalling and contributes to stabilize BMP signalling on the dorsal side via a positive feedback circuit (Lapraz et al., 2009). In addition, the BMP ligands Admp1 and Admp2 damp BMP signalling fluctuations through an expansion-repression mechanism (Ben-Zvi et al., 2008; De Robertis, 2009; Joubin and Stern, 2001; Kimelman and Pyati, 2005; Lapraz et al., 2015; Lele et al., 2001; Reversade and De Robertis, 2005; Willot et al., 2002). This mechanism, which relies on the transcriptional repression of *admp1* expression by BMP2/4 signalling, allows *admp1* expression to increase and ADMP1 protein to be shuttled to the dorsal side by Chordin when BMP signalling decreases. Thus, an increase in *admp1* expression compensates for the reduction of the intensity of BMP signalling.

One prominent feature of the D/V onset specification is that it relies extensively on the maternal inputs Panda and Univin, which respectively represses and promotes ventral fate (Range et al., 2007, Haillot et al., 2015, reviewed in Molina and Lepage, 2020). Univin is a TGF-β related to Vg1 and GDF1/3. Univin is essential for Nodal signalling through the Nodal/Activin receptors; in absence of Univin, Nodal expression stops and D/V axis specification fails. Panda is a member of the TGF-β superfamily presumed to repress ventral fate by a still unidentified mechanism (Haillot et al., 2015). Finally, the transcriptional repressor Yan/Tel acts as a negative regulator of *nodal* expression, whose expression is required downstream of Panda (Molina et al., 2018).

Previous studies have shown that Nodal produced by the ventral ectoderm is a strong ventralising signal (Duboc et al., 2004; Lapraz et al., 2015; Saudemont et al., 2010). Overexpression of *nodal* causes all ectodermal cells to adopt a ventral fate (Fig. 1B), whereas a loss of Nodal function prevents specification of both ventral and dorsal fates and causes the ectoderm to differentiate as a ciliary band (Fig. 1C). Conversely, the activity of BMP2/4 protein promotes dorsalisation, and overexpression of BMP2/4 forces all ectoderm cells to adopt a dorsal fate (Haillot et al., 2015; Lapraz et al., 2009; Lapraz et al., 2015) (Fig. 1D). In contrast, removing the function of the BMP2/4 ligand from fertilisation on prevents specification of dorsal fates, leading to formation of an ectopic ciliary band territory in the dorsal region (Fig. 1E). Additionally, knocking down *panda* expression in this BMP2/4 loss-of-function experiment enables *nodal* to be expressed through the dorsal side of the ectoderm and promotes ventral fates in all ectodermal cells (Fig. 1F). Conversely, a local *panda* overexpression promotes dorsal fates, suggesting that *panda* is sufficient to break the radial symmetry of the embryo and necessary to specify the D/V axis (Haillot et al., 2015). The BMP and Nodal ligands thus show strongly antagonistic activities. However, the mechanism underlying this antagonism and the resulting cell fate decision still awaits clarification.

Due to the largely non-cell autonomous nature of the D/V GRN, to the many events of protein diffusion, and to the intertwined feedback circuits involved, an intuitive understanding of the logic of the network is hard to obtain. For example, Nodal and BMP2/4 are co-expressed in the ventral territory, but the corresponding signalling pathways are operating at opposite poles of the D/V axis. In this context, a model of the D/V gene regulatory network is very useful to formalise the complex regulatory interactions at stake (Wilczynski and Furlong, 2010). A Gene Regulatory Network can be modelled as a static regulatory graph using standardised graphical conventions to represent the molecular interactions between relevant regulator components (Arnone and Davidson, 1997). Such regulatory graphs can be supplemented with threshold levels and regulatory rules to obtain a predictive, dynamical model (Mbodj et al., 2016; Peter et al., 2012; Thomas and D’Ari, 1990).

In the present study, we built a logical model (i.e. using Boolean algebra for the regulatory rules) of the sea urchin dorsal-ventral specification GRN to (i) assess its accuracy, (ii) compare the simulations of different perturbations with the observed gene expression patterns, (iii) explore the dynamics of the system, and (iv) develop a multicellular framework to test the ability of the model to generate the spatial patterns observed in wild type and perturbed embryos.

## Results

### Model building

We constructed a model of the GRN driving the D/V patterning of the sea urchin ectoderm. We started by compiling experimental data to identify the key genes and regulatory interactions (Fig. 2). The raw data that provided the spatial and temporal expression information to build the model were derived from the analysis of high resolution in situ hybridizations, Northern blot and systematic perturbations experiments. Loss-of-function experiments via morpholino injections are particularly important for GRN reconstruction since they allow to test if a gene is required for activation of another gene. Gain-of-function experiments via mRNA injection were also used in many instances to test for the ability of a given gene to induce another gene when overexpressed.

**Figure 2.**
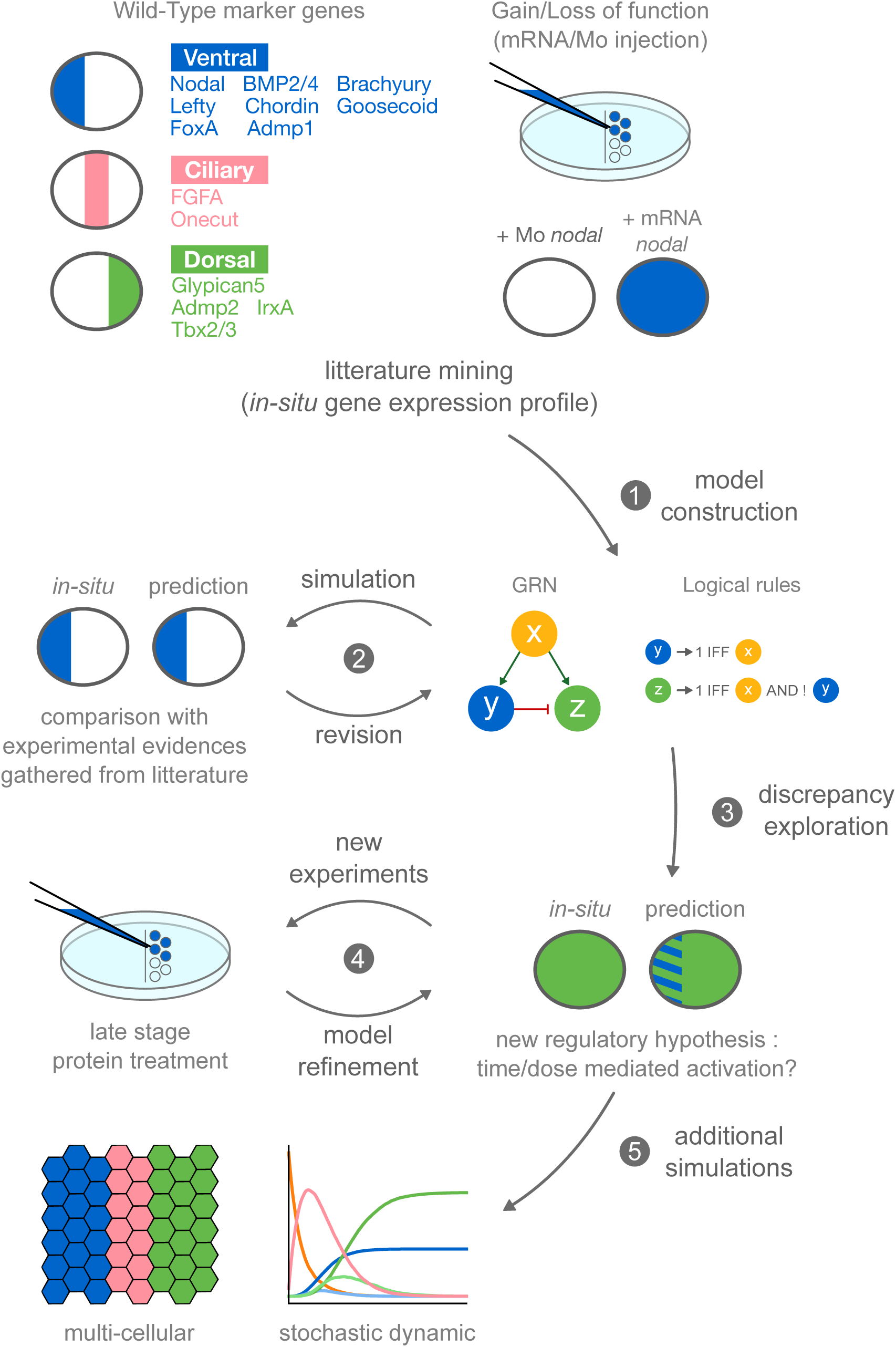
Iterative integration of biological data into the GRN model. The GRN model has been built through an iterative process. A first version based on literature curation and experimental evidence was set and then simulated in wild type and perturbation conditions. The results of wild-type and morphant simulations were systematically compared with experimental results. In case of discrepancy, the regulatory graph and the logical rules were refined, and the behaviour of the model was then re- examined through the same process (1-2). Following this iterative process, the unsolved discrepancies were explored by novel experiments designed to test the different regulatory hypotheses of the model (3-4). The final model was then further extended and analysed using additional tools (MaBoSS for stochastic simulations, EpiLog for multicellular simulations).

Based on these data, we first built a regulatory graph representing the D/V GRN in a single cell, using the software GINsim (ginsim.org) (Naldi et al., 2018). The model has been built through an iterative process, alternating simulations and model refinements, to optimise our simulation results as compared to experimental observations (Fig. 2). An extensive analysis of published data enables us to delineate relevant regulatory interactions and formalise them in terms of a signed, directed graph, complemented with logical rules (see Material and methods). The resulting model was then subjected to computational analyses, which were confronted to experimental data. The analysis of discrepancies led us to revise the network and the regulatory rules (e.g. by incorporating a new interaction), until the simulations qualitatively recapitulate all relevant experimental observations (Fig. 2). The main challenge encountered during this process consisted in the integration of multiple interactions targeting the same component. Indeed, the number of possible regulatory rules exponentially increases with the number of incoming interactions. The regulatory rules must correctly reflect the relative importance of each incoming signal, taking into account different qualitative ranges. These rules are encoded using the classical logical operators (AND/OR/NOT), together with multilevel variables when justified (see Material and methods).

At the end of this iterative process, the model was further refined based on the results of novel experiments designed to disentangle the interplay between Nodal and BMP2/4 pathways (Fig. 2).

Regarding the computational analyses performed, we combined three main approaches: i) a direct computation of stable states for relevant input combinations, ii) stochastic simulations to estimate the probabilities of alternative cell fates (*chordin* morpholino), and iii) logical multicellular simulations to cover intercellular signalling events (see Results).

We present hereafter the final model and selected simulations recapitulating the patterns observed experimentally. The model encompasses a total of 30 nodes, linked by 23 activations and 15 inhibitions (Fig. 3). The nodes included into the model correspond to signalling and regulatory components, while the signed arcs denote regulatory interactions between these components. Signalling factors are modelled as input nodes (in yellow in Fig. 3), providing activatory or inhibitory signals through the corresponding membrane receptors.

**Figure 3.**
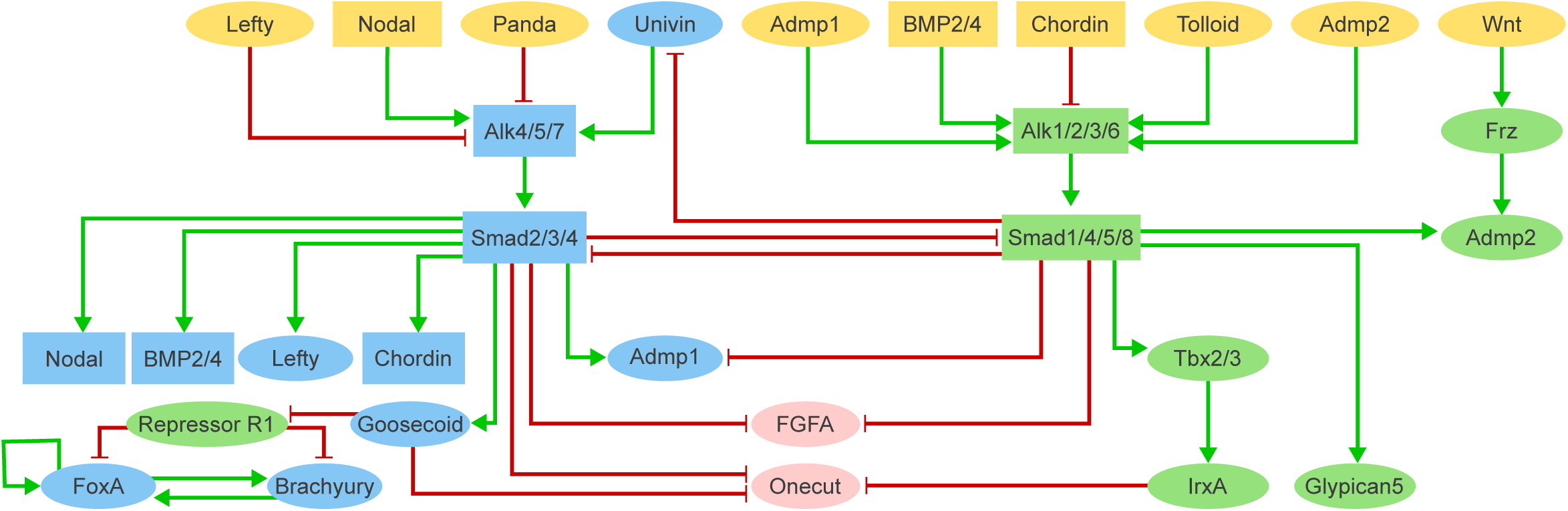
Logical model integrating the main signalling pathways controlling the early dorsal-ventral axis specification in the sea urchin *Paracentrotus lividus* embryo. Relying on a logical formalism, this model was defined and analysed using the software GINsim. Green and red arrows represent activations and repressions, respectively. Ellipsoid and rectangular components represent Boolean and multivalued components, respectively. The components in yellow correspond to model inputs. Internal components are coloured according to their domain of expression along the dorsal-ventral axis, i.e. dorsal (green), ventral (blue) or ciliary (pink) regions.

Each (non-input) component is classified as ventral (eleven nodes shown in blue in Fig. 3), ciliary band (two nodes shown in pink in Fig. 3) or dorsal (eight nodes shown in green in Fig. 3), according to the reported location and time of activation in the presumptive ectodermal regions. For example, *goosecoid* is activated by the Nodal cascade in the ventral region in wild-type condition, and is thus considered as a ventral gene. The model encompasses the main regulatory components of TGF-β signalling pathways, including the ligands, negative regulators such as the proteins that trigger receptor degradation, downstream transcription factors, and antagonists. Each component of the model is annotated with textual explanations and database links (in particular to PubMed) documenting our modelling assumptions (main references include the GRN diagram published for *Paracentrotus lividus* in Haillot et al., 2015; Lapraz et al., 2006; Lapraz et al., 2015; Molina et al., 2013; Range et al., 2007; Saudemont et al., 2010).

Among the 30 components of the model, 20 are associated with Boolean variables (ellipsoid nodes in Fig. 3, taking the values 0 or 1 depending on their activation state), while the remaining components are associated with multilevel variables (rectangular nodes in Fig. 3, associated with three or four integer levels, from zero to 2 or 3, see below). The nine input nodes (shown in yellow in Fig. 3) define 576 possible configurations of input values. Using the Java library bioLQM (Naldi, 2018), we identified 654 stable states, which can be split into three main patterns based on the active nodes: 288 ventral, 126 ciliary and 240 dorsal patterns (cf. Jupyter notebook).

### An antagonism between the Nodal and BMP2/4 pathways drives the allocation of cell fates along the dorsal-ventral axis

A key feature of our model consists in the antagonistic activities of BMP2/4 and Nodal. In this respect, additional experiments were conducted to probe the underlying mechanisms. We first tested whether difference in dosage between the corresponding signalling molecules could favour the establishment of a specific cell fate. Treatment with an intermediate dose of BMP2/4 protein resulted in embryos developing with a straight archenteron, no mouth, and covered with a ciliary band like ectoderm (Fig. 4A), which is a prototypical *nodal* loss-of-function phenotype. In contrast, for intermediate doses of Nodal, the embryos developed with a reduced apex, reminiscent of the BMP2/4 loss-of-function phenotype. These observations suggest that ectodermal cells receive both antagonistic ventralising Nodal and dorsalising BMP2/4 signals, and integrate them even at intermediary doses at the level of the *cis*- regulatory sequences of their target genes.

**Figure 4.**
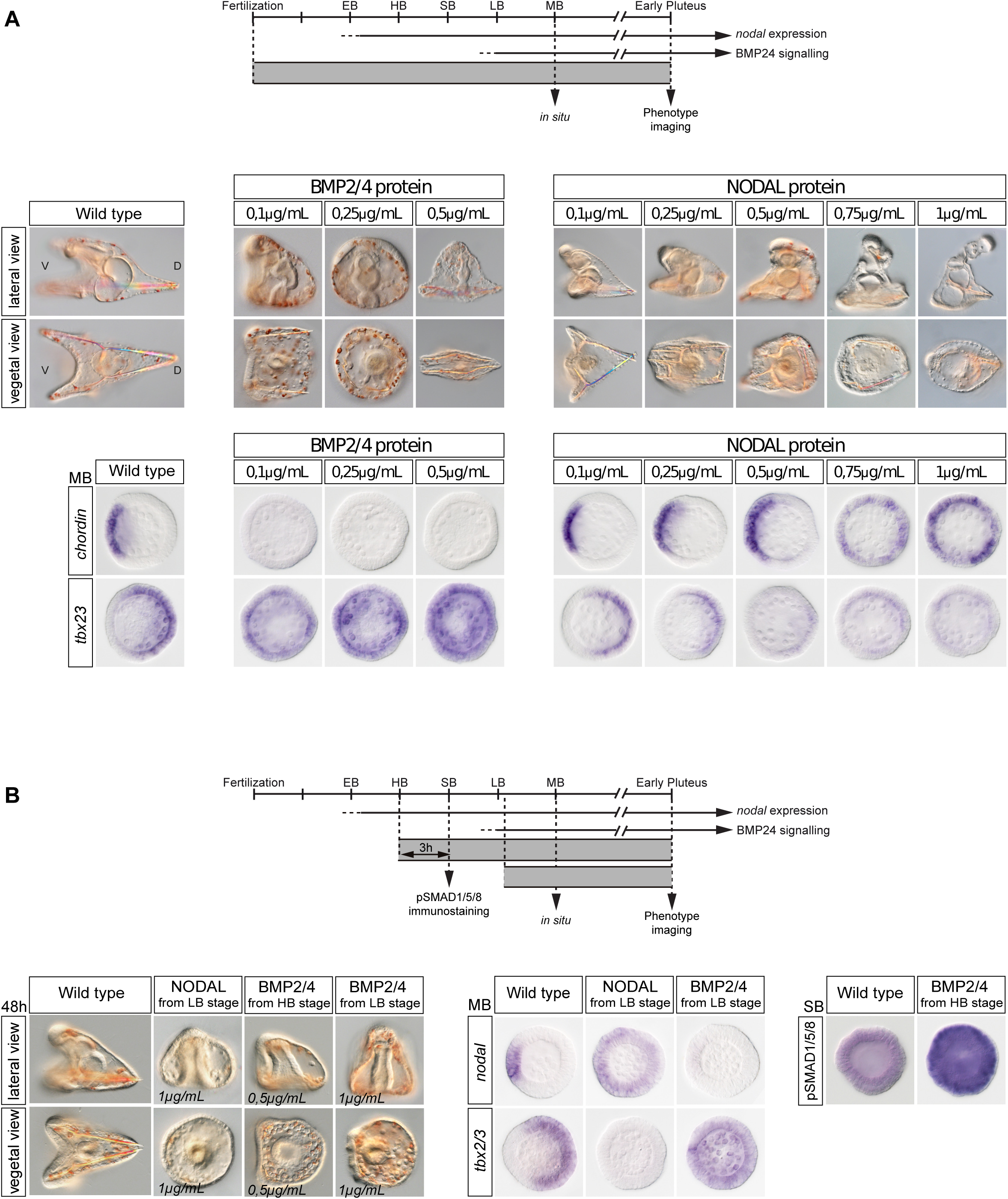
BMP and Nodal signalling antagonize each other to pattern the D/V axis of the sea urchin embryo. (A) Continuous Nodal and BMP2/4 protein treatments at increasing concentrations from the fertilized egg stage. Increasing concentrations of BMP2/4 protein treatment progressively dorsalises the embryo at expenses of the ventral territories, as reflected by the expansion of the expression of the dorsal marker *tbx2/3* and the repression of the expression of the ventral marker *chordin*. On the other hand, increasing concentrations of Nodal protein treatment gradually ventralises the embryo at the expenses of the dorsal territories as reflected by the gradual expansion of the expression of the ventral marker *chordin* and the repression of the expression of the dorsal marker *tbx2/3*. (B) Three hours Nodal or BMP2/4 protein treatments at late blastula and hatching blastula stages are sufficient to cross-antagonise each other signalling. Three hours Nodal protein treatment at late blastula stage results in rounded-shaped embryos partially ventralised that overexpress the ventral marker *nodal* at the expenses of the dorsal marker *tbx2/3*. Complementary, three hours BMP2/4 protein treatment at hatching or late blastula stages promotes massive pSMAD1/5/8 signalling and results in partially dorsalised embryos that overexpress the dorsal marker *tbx2/3* at the expenses of the ventral marker *nodal*. EB, Early Blastula; HB, Hatching Blastula; SB, Swimming Blastula; LB, Late Blastula; MB, Mesenchyme Blastula; V, Ventral; D, Dorsal.

However, since these treatments were performed soon after fertilisation, it was not clear whether the outcome was reflecting an antagonism between Nodal and BMP occurring during cell fates allocation, or if it resulted from one pathway being activated early and dominantly in all cells of the embryo following the injection of mRNA into the egg. To address this issue, we repeated the Nodal and BMP2/4 treatment at late blastula and at early mesenchyme blastula stage. Treatments with Nodal or with BMP2/4 proteins at late blastula radialised the embryos by respectively inducing ventral or dorsal fate in all ectodermal cells (Fig. 4B). These results confirm that Nodal and BMP2/4 pathways act antagonistically also during fate specification and that a dosage competition plays an important role in cell fate specification.

These results led us to pay a particular attention to the encoding of this antagonism in our model. First, to account for dosage effects, we associated ten nodes of the model with ternary variables (rectangular nodes in Fig. 3, taking values from 0 to 2). These multivalued nodes allow for a more fine-grained encoding of the activation states of key morphogens and downstream components whose effects are dose-sensitive (nodes Nodal, Chordin, BMP2/4, Alk4/5/7, Alk2/3/6, Smad2/3/4, Smad1/4/5/8 in Fig. 3). For example, Nodal activity can either overcome (level 2) or be counteracted (level 1) by the inhibition of Lefty. Additionally, Nodal input node is represented by a quaternary variable, in order to distinguish expression levels from wild-type (level 1-2) versus gain-of-function (level 3) conditions. Second, the antagonism between the Nodal and BMP pathways is encoded as a reciprocal inhibition between Smad2/3/4 and Smad1/4/5/8 (Fig. 3), which implements the competition of these signalling complexes for the shared molecule Smad4. Noteworthy, each of these two inhibitory interactions can be counteracted by an increased activity of the other antagonistic pathway, according to our dose-dependent competition hypothesis (this is encoded in the corresponding logical rules, see Table 1).

**Table 1.**
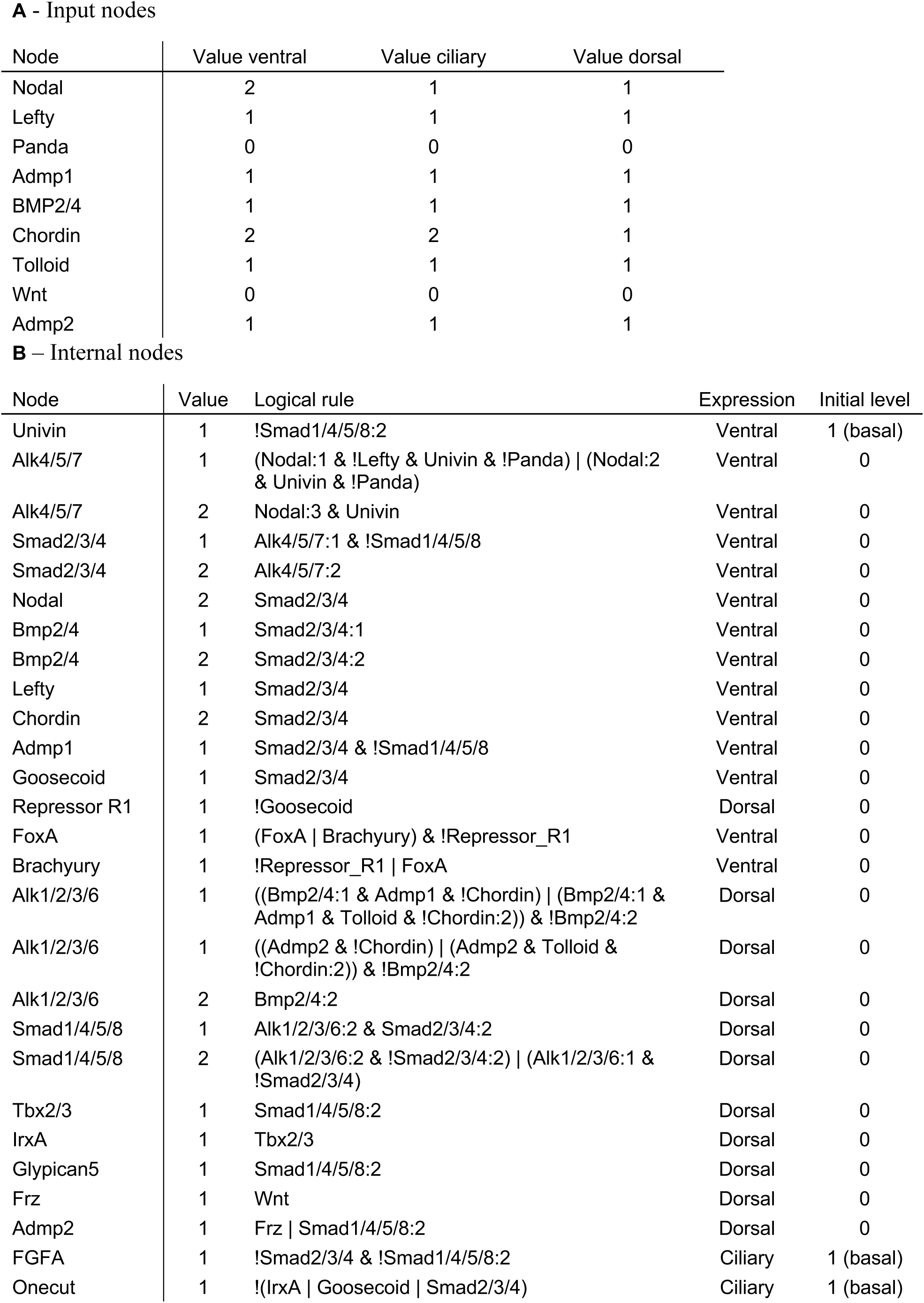
Logical rules of the unicellular model. Logical rules are used to define the behaviour of each node, for each territory, relative to its direct upstream regulatory nodes. Input nodes (A) do not have any assigned rules, as they are set to a given fixed value specific for each territory when performing simulations (value shown for late stage simulation). For the internal nodes (B), the table lists the logical rule required to reach non zero levels, a as well as the wild-type initial level. All nodes have their basal level set to 0 (inactive), excepting Univin, FoxA and FGFA (basal level 1), as they tend to be ubiquitously active without the need of explicit activators. The logical rules combine literals, each representing the activity of one node, with the Boolean operators OR (“|”), AND (“&”) and NOT (“!”).

### Model stable states match experimental wild-type and morphant phenotypes

To test our model, we ran simulations and compared the resulting stable states with the wild type pattern observed experimentally. We applied different sets of values for the input nodes, each set corresponding to a specific territory of the ectoderm (Table 1 and materials and methods). For the relevant combinations of active inputs, the resulting stable states are compared with the list of marker genes expected to be expressed in the corresponding territory, based on *in-situ* hybridization experiments. We first considered the combinations of inputs corresponding to early 32-cell stage embryos signalling (Fig. 5), which corresponds to the onset of specification of the ventral organiser, forming at the opposite side of the gradient of Panda mRNA (Haillot et al., 2015). This pattern is correctly recapitulated by the stable states obtained for the wild-type situation (Fig. 5). The resulting stable states were then used to specify the input conditions reflecting a later blastula stage, after diffusion and shuttling of zygotic factors have taken place (Fig. 5). Indeed, as multiple diffusion events occur, some model inputs expressed in one territory are active in a broader region for blastula simulations (Fig. 6). After some iterative refinements of the rules, model simulations qualitatively recapitulated the expression patterns expected for each individual territory (ventral, ciliary, dorsal) (Fig. 6). Hence, we can conclude that, the regulatory graph shown in Fig. 3 supplemented by the logical rules of Table 1 are sufficient to specify the three main ectodermal D/V patterns of the sea urchin embryo.

**Figure 5.**
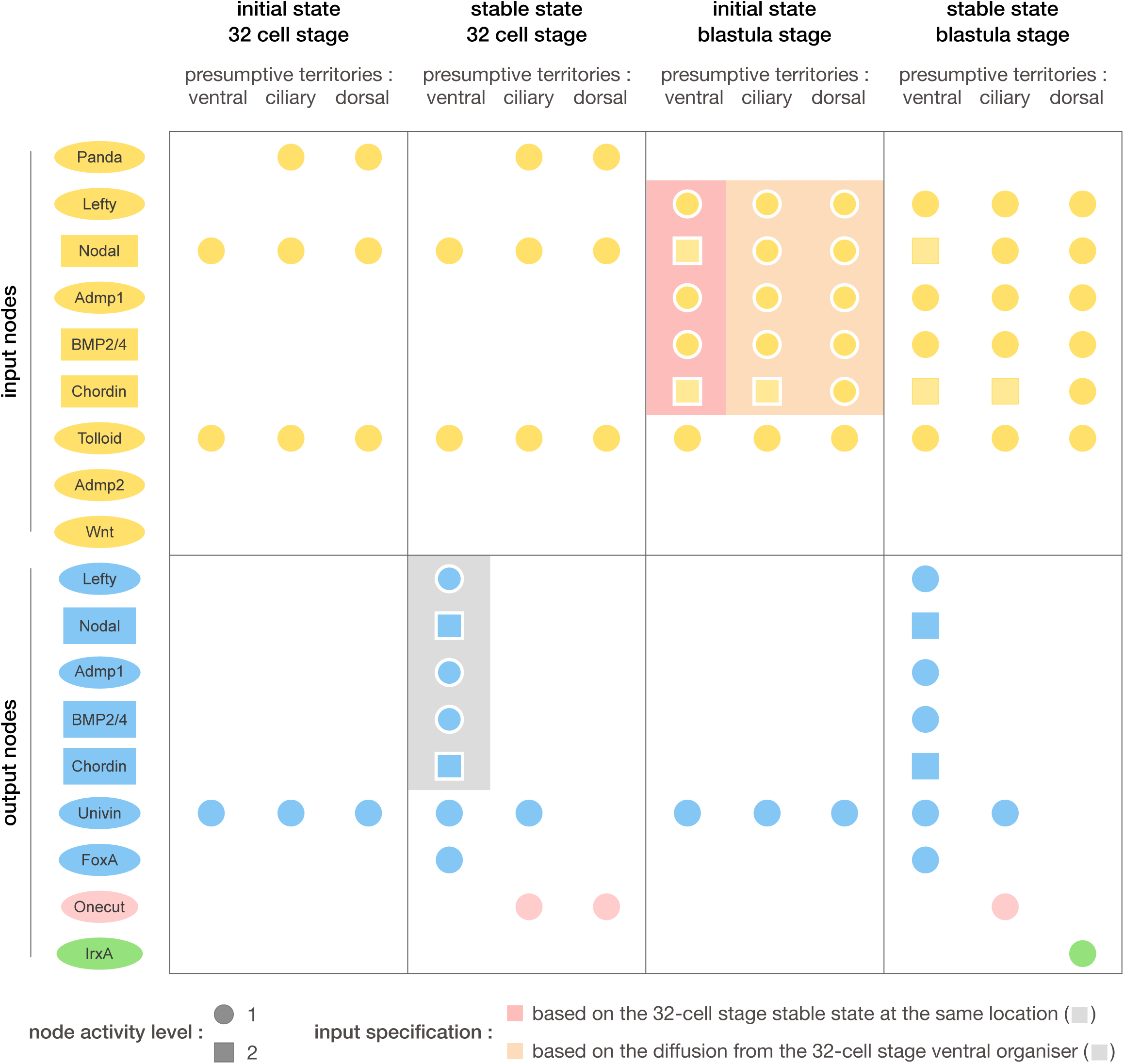
Simulation of early 32-cell stage and specification of later stage inputs. By restricting the active input nodes to combinations of Nodal, Panda and Tolloid (1st column), our unicellular model recapitulates the wild-type patterns observed in the 32-cell stage embryos (2nd column), where *panda* is expressed in the presumptive dorsal region and restricts *nodal* expression in the presumptive ventral region (Haillot et al., 2015). The output node values resulting from our ventral wild-type simulation (grey background) were then used to define the input node values for the ventral simulation of later developmental stages (red background). Additionally, taking into account the diffusion and shuttling events known to occur from the ventral region to further dorsal territories in this developmental time window, the output values of the 32-cell stage ventral simulation were also used to define the input node values for the simulation of the ciliary and dorsal regions in the blastula stage simulation (orange background).

**Figure 6.**
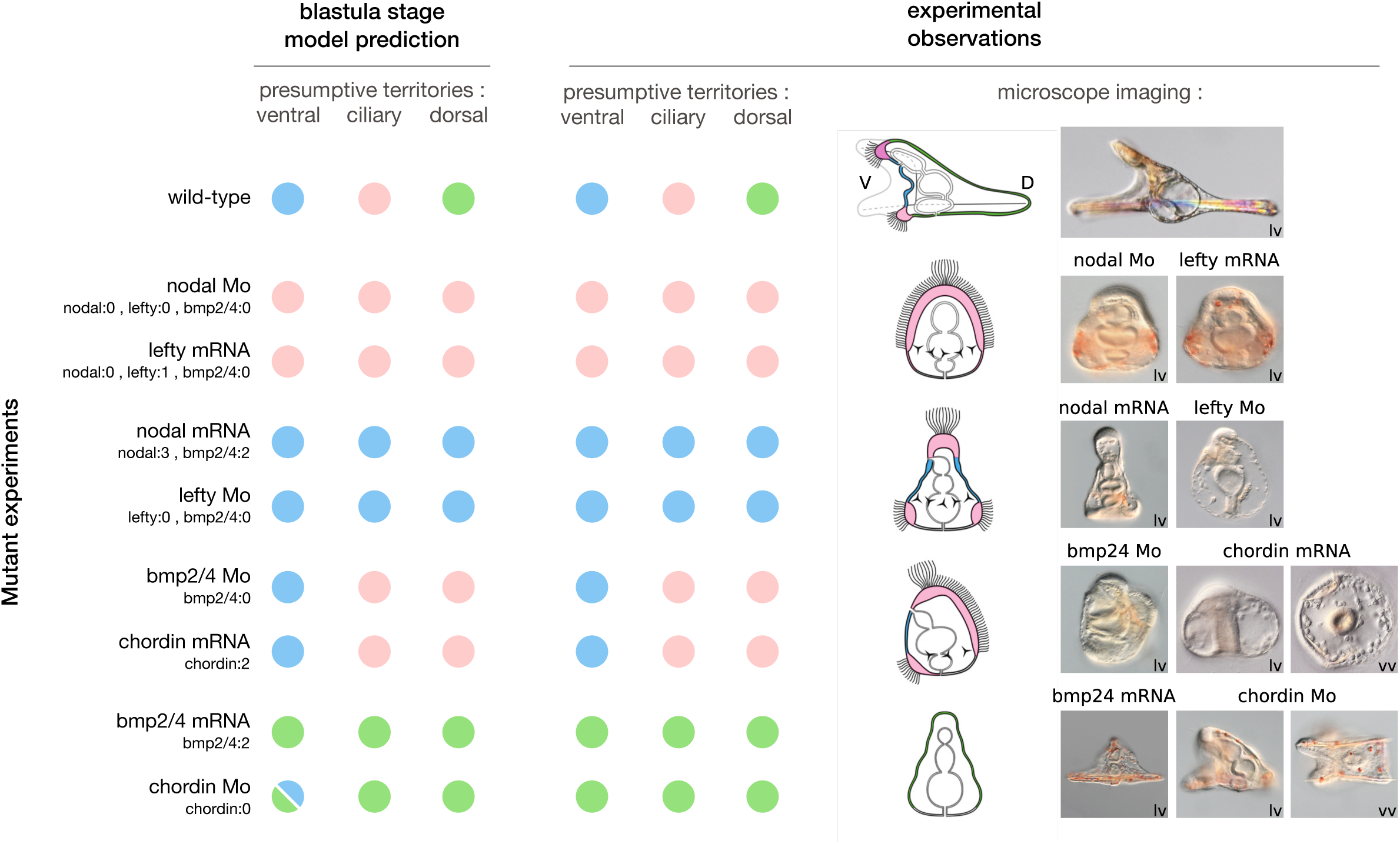
Comparison of blastula model simulations and experimental results. With proper logical rules (see Table 1), inputs and initial state conditions (see Fig. 5), our model gives rise to different stable patterns (circles, left), which qualitatively match experimental observations of dorsal (green), ventral (blue) and ciliary (pink) territories (circles, schematic and microscope imaging, right). Note that, in the case of the *chordin* morpholino, the model predicts two possible stable patterns in the ventral region, whereas experiments point to a weak ventral specification phenotype.

To further validate and explore the properties of our model, we simulated loss- and gain-of-function experiments (mRNA or morpholino injection) by restricting the range of reachable levels for one or multiple node(s), e.g. to zero for a loss-of-function, or to a higher value for a gain-of-function (cf. material and methods). As in the wild-type conditions, we challenged our model by comparing the resulting stable states with the *in-situ* patterns observed following mRNA or Morpholino injection in the embryo at early stage. For seven of the eight morphants simulated, the model returned a unique single stable state in each region, which qualitatively matched experimental observations (Fig. 6).

In the cases of *nodal* Morphants and of *lefty* mRNA overexpression, the ventral cascade fails to be established, leading to the absence of both Nodal and BMP2/4 pathway activities, and to the presence of a default ciliary state in all the ectodermal cells. Following *nodal* mRNA overexpression, the competition between Smad2/3 and Smad1/5/8 for Smad4 turns in the advantage of the ventral cascade producing a fully ventralised embryo. The same pattern is obtained for the *lefty* Morpholino, because this perturbation impacts the diffusion-repression mechanism controlled by Lefty (Juan and Hamada, 2001), enabling *nodal* expression to propagate without restriction (Duboc et al., 2008). In the case of overexpression of the dorsal fate repressor *chordin,* or yet in the case of BMP morphants, the absence of BMP2/4 signalling fosters a ciliary band state in the presumptive dorsal territory. However, as BMP2/4 is not necessary for the expression of *nodal*, the ventral cascade maintains a wild type expression pattern in these morphants. Finally, following *bmp2/4* overexpression, the competition for the common Smad is driven toward the activation of the dorsal cascade, giving a fully dorsalised ectoderm state.

Interestingly, in the case of the *chordin* morpholino, the model returned two stable states (denoted by two blue and green half circles at the bottom and left of Fig. 6) in the presumptive ventral region, corresponding to ventral and dorsal fates, respectively. This situation is further investigated using a probabilistic framework hereafter.

### Stochastic logical simulation of the chordin Morpholino

Using a probabilistic extension of the logical framework, one can unfold the temporal dynamics of the regulatory network for given initial conditions, and estimate the prevalence of a given cellular fate when the model predicts alternative stable states. In the case of the ventral region in *chordin* morphant conditions, we have seen that our model predicts two different stable states, corresponding to ventral and dorsal expression patterns. Using the software MaBoSS (https://maboss.curie.fr) (Stoll et al., 2017), we performed stochastic temporal simulations of our model to generate mean time plots and estimate the probability to reach each of these stable states. In the absence of precise kinetic information, we first used equal rates for all (up or down) transitions.

In the wild type ventral region, as expected, all stochastic simulations gave rise to a ventral expression pattern (Fig. 7A). In contrast, for the *chordin* morphant, the dorsal state is reached about twice as often as the ventral state (Fig. 7B). In other words, the dorsal pathway is more likely to win the competition for Smad4. This partial dominance of the dorsal pathway matches the weak dorsal patterns observed experimentally for *chordin* morphants (Fig. 6), which presumably result from the co-activation of the two antagonistic pathways. Additionally, the prediction of a transient expression of ciliary genes in the ventral region (Fig.7) is further supported by experimental evidence from Saudemont et al. (2010).

**Figure 7.**
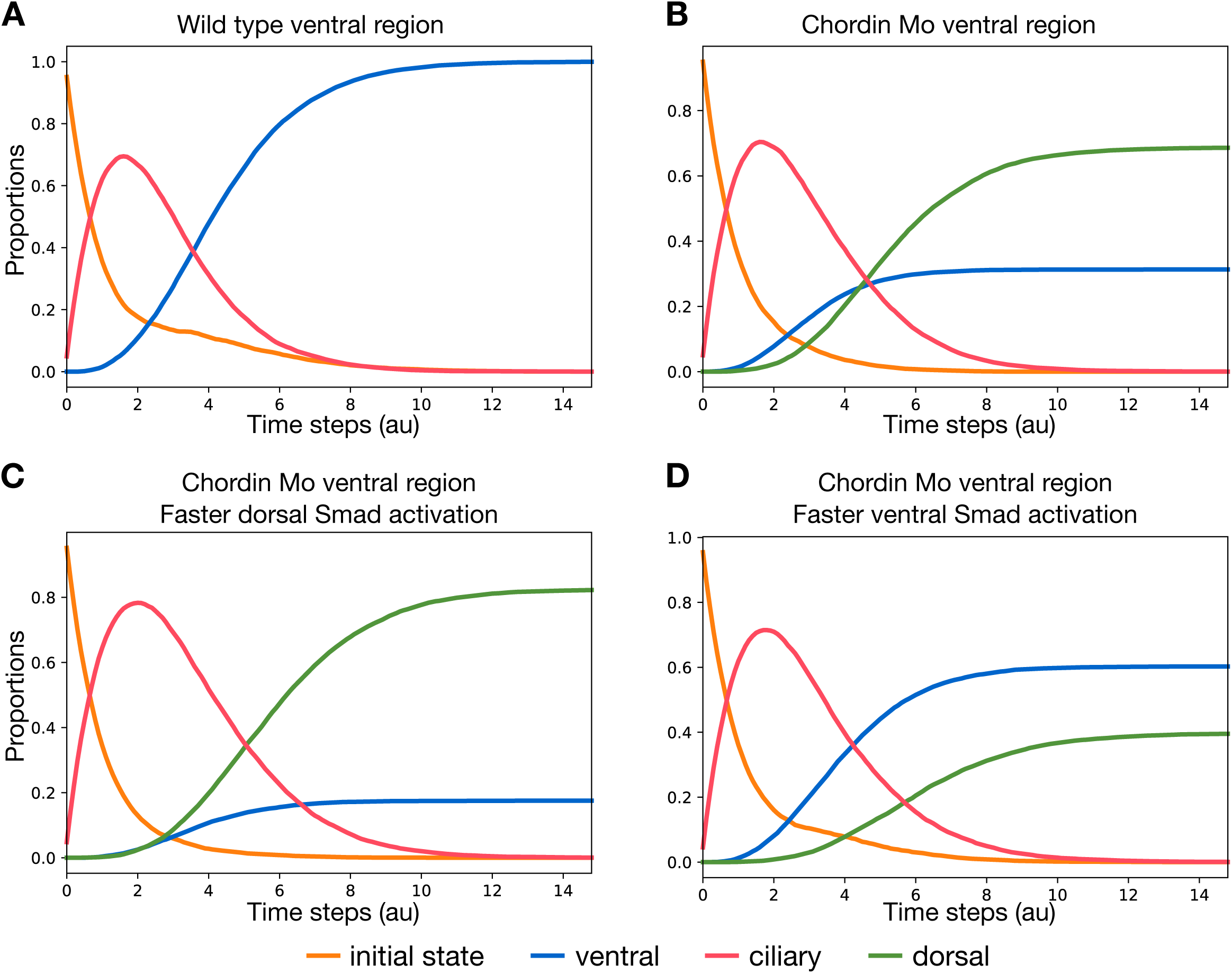
Probabilistic time-course simulations of the unicellular model starting with the ventral initial state. These plots show the temporal evolutions of the mean activation levels of Goosecoid, Iroquois and Onecut, representing ventral (blue), dorsal (green) and ciliary (pink) phenotypes, respectively. All simulations start from a ventral initial state (orange). The first plot (A) corresponds to the wild type, while the three other ones (B, C, D) correspond to *chordin* knock down conditions. Simulations (A, B) were performed with equal up and down state transition rates. Further simulations were performed using rates favouring the dorsal cascade (C), or favouring the ventral cascade (D) (see supplementary material and methods for details).

In the chordin morphants, BMP signalling diffuses unrestricted in the ventral ectoderm, promoting dorsal fates and repressing the ventral cascade. However, since Nodal is critically required for *bmp2/4* activation, *nodal* down-regulation in turn leads to the repression of BMP2/4 signalling. Therefore, in the absence of Chordin, both the ventral and dorsal cascades are activated. This conclusion is supported by the experimental observation of transient Smad1/5/8 signalling and *tbx2/3* expression in the ventral ectoderm (Lapraz et al., 2009).

To further assess whether this imbalance in favour of the dorsal pathway activation is sensitive to kinetic (transition) rates, we ran stochastic simulations with different Smad activation rates for the two pathways. An imbalance in favour of the dorsal Smad activation further increases the final proportion of dorsal fate compared to wild type (Fig. 7C). On the contrary, an imbalance in favour of the ventral Smad activation inverses the relationship, with a higher fraction of ventrally specified states compared to dorsally specified states, almost mirroring the ratios obtained with equiprobable transition rates (Fig. 7D).

In conclusion, in the condition of a simultaneous activation of both ventral and dorsal cascades without inhibition (i.e. *chordin* morphants), the outcome of the competition between the BMP2/4 and Nodal pathways is at least partly driven by the kinetic rates of Smad activation. In such situation, the pathway-specific Smad firstly activated reaches more quickly a concentration level sufficient to repress the other pathway by pre-empting Smad4 and thereby fosters the corresponding state stable.

### Multicellular simulations emphasise the crucial role of long-range signal diffusion

In the preceding section, we simulated the behaviour of cells for each of the three different presumptive territories by selecting appropriate combinations of signalling input levels, which were considered as fixed during the whole duration of the simulations. To model more precisely the production and diffusion of signalling molecules across the ectoderm, we used the software EpiLog (https://epilog-tool.org) (Varela et al., 2019), which supports simulations of an epithelium encompassing multiple cells connected by diffusing signals. The behaviour of each cell of the epithelium is modelled by the same cellular logical model, but levels of input signals directly depend on the output signal values from neighbouring cells. The input signals perceived by a given cell are integrated into logical diffusion rules, which are updated synchronously during simulations (see Materials and methods and Table 2) (Fig. 8A).

**Figure 8.**
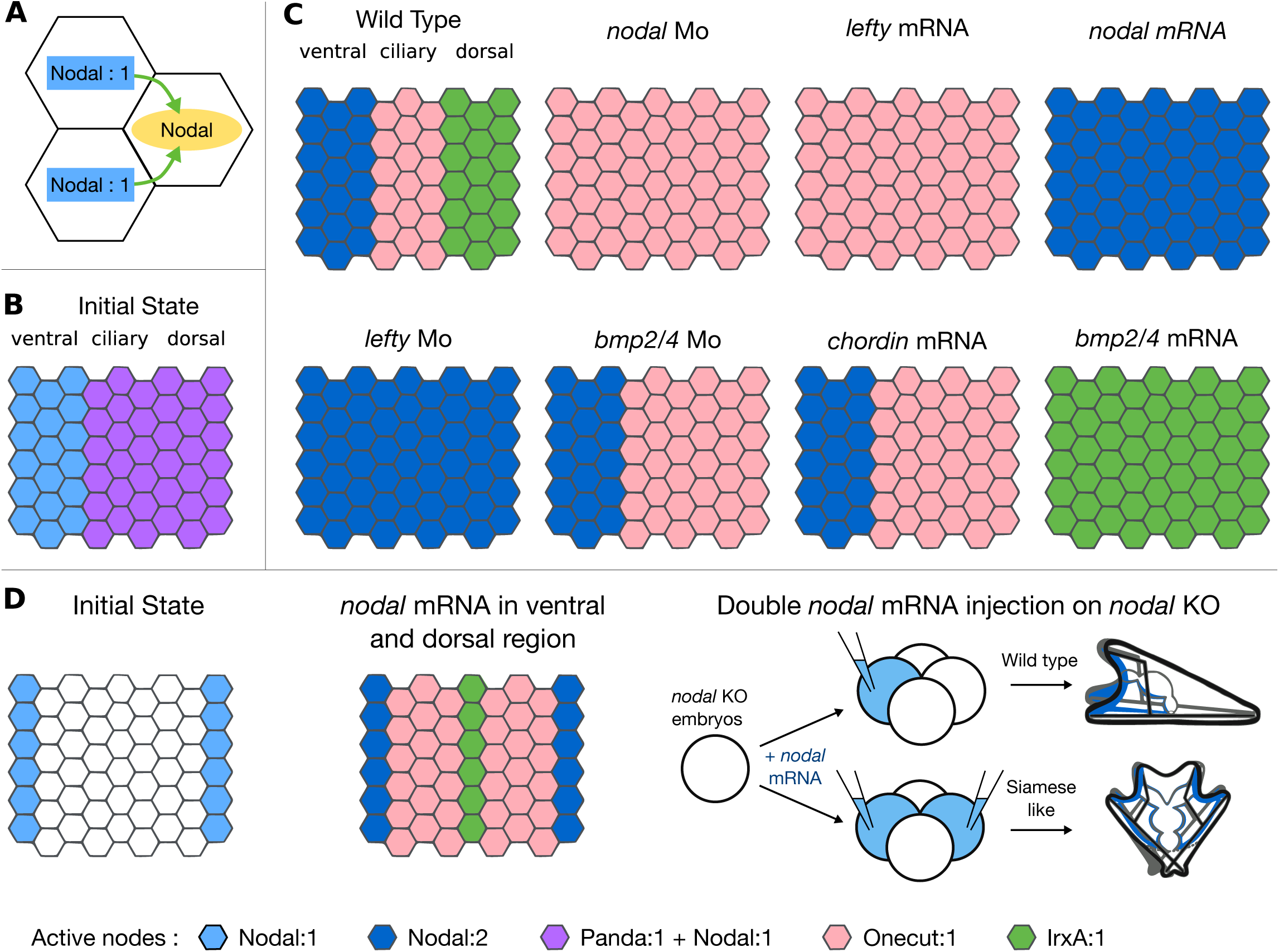
Multicellular logical simulations for wild-type and morphant conditions, using the software Epilog. Across the multicellular epithelium, specific logical rules have been defined to model the diffusion of signalling components (A) (see Table 2 for diffusion rules). At the initial state, Panda is presents in the presumptive ciliary and dorsal territories, while Nodal is ubiquitously expressed in the whole epithelium (B). Multicellular simulation results for the wild-type and morphant conditions (C) qualitatively match our experimental results. We further simulated the injection of Nodal mRNA in a four-cell embryos in two opposite cells by initiating the model with two regions expressing *nodal* at the opposite poles of the epithelium (D, 1st and 9th columns) which resulted in a morphant displaying a symmetric pattern of ventral, ciliary and dorsal territories along the dorsal-ventral axis, as observed by Lapraz et al., 2015 (D).

**Table 2.**
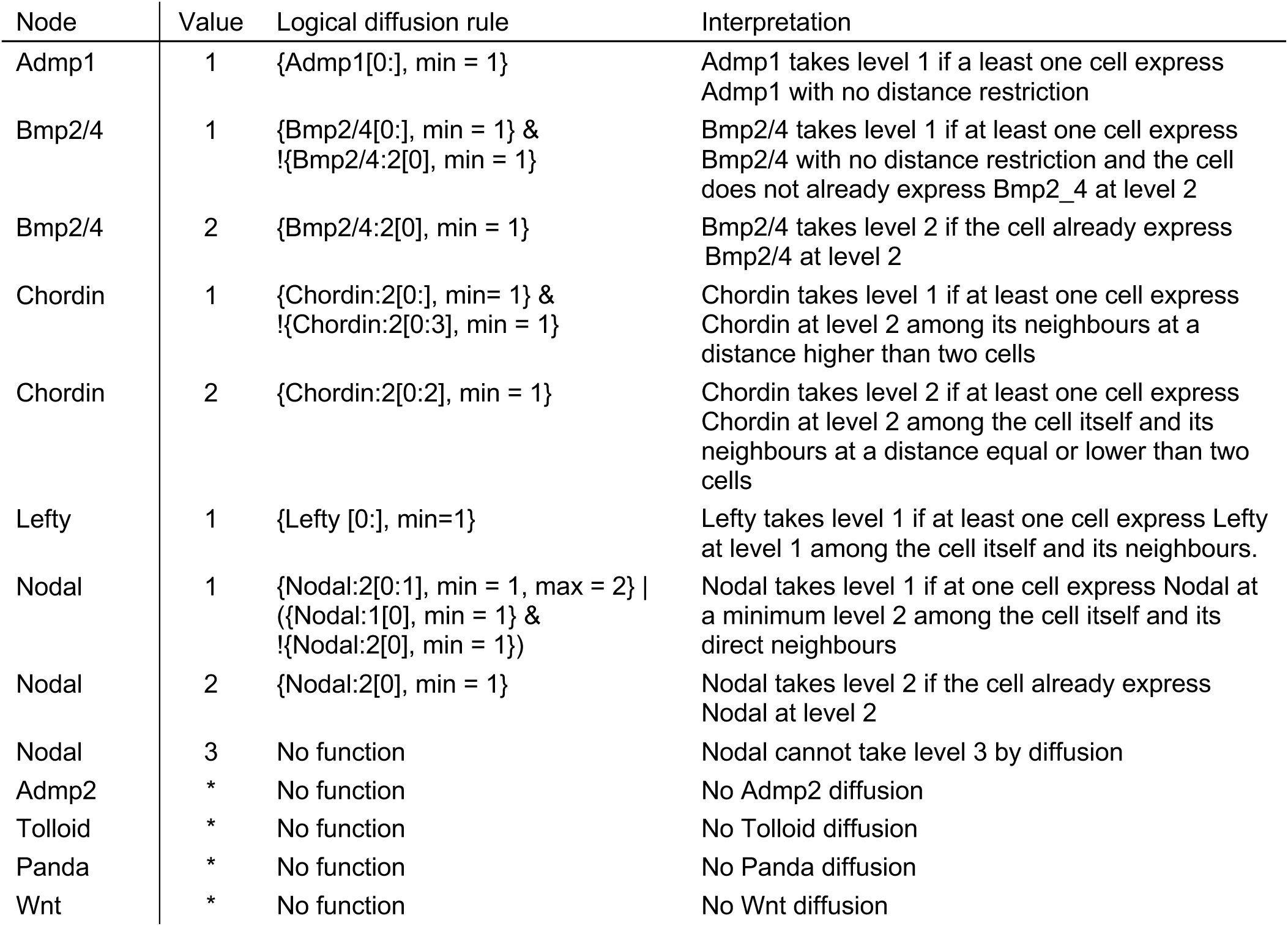
Logical diffusion rules used in Epilog. Logical rules are used to define the diffusion dynamics perceived by the input nodes, depending on the values of the output nodes in the neighbouring cells. Diffusion rules are defined in the format “{N:L[D],S}”, with N as the node with perceived diffusing signal, L its required activation level, D the distance range to perceive diffusion and S the minimum and/or maximum number of cell required in this state. For example, the fifth row specifies that cells will have their Chordin input node value converging toward the value 2 if at least 2 cells are expressing Chordin at a value 2 at a maximum distance of three cell.

To simulate the wild type situation, we initialised the model with a broad weak expression of Nodal and Panda expression in the presumptive ciliary and dorsal territories (corresponding to the six right-most range of cells), as observed experimentally (Haillot et al., 2015) (Fig. 8B). The simulation correctly recapitulates the expected contiguous ventral, ciliary and dorsal ectoderm territories (Fig. 8C). This result suggests that the relatively simple diffusion rules properly account for the dynamics of the proteins governing the dorsal-ventral patterning. In addition, this result highlights the crucial role of Nodal to direct specification of the ectoderm along the whole dorsal-ventral axis. Interestingly, in the course of wild-type simulation, we can clearly distinguish the main steps of the D-V patterning, with the initial restriction of Nodal expression on the ventral side, followed by the diffusion of BMP2/4 signalling toward the dorsal side, until it activates the dorsal marker genes (see Fig. S1).

As in the case of the unicellular model, we can apply specific perturbations to assess their impact at the tissue level. As shown in Fig. 8C, our multicellular simulations accurately recapitulate the phenotypes of the different morphant patterns observed experimentally. Note that the chordin morpholino has been discarded from these simulations, as it gave rise to two different stable states, which cannot be properly simulated with the Epilog deterministic input updating approach.

Using Epilog, it is further possible to perform local perturbations by modifying the initial levels of one or several signalling molecules at specific epithelium locations. Using this feature, we could recapitulate in silico the results of an experiment reported by (Lapraz et al., 2015), who injected *nodal* mRNA into two opposite cells of a 4-cell stage *nodal* knock-down embryo (i.e. following a *nodal* Morpholino injection in the egg). This experiment triggered the formation of an ectopic, inverted D/V axis and resulted in the development of siamese pluteus larvae with two ventral sides, two ciliary bands and a central dorsal territory. Using Epilog and imposing *nodal* and *smad2/3/4* activity at the initial state in both the ventral and the dorsal side of the epithelium, our spatial logical simulation qualitatively recapitulates the siamese pattern observed experimentally (Fig. 8D).

## Discussion

Gene regulatory networks integrate documented interactions between transcription factors, signalling components, and their target genes, which ultimately translate the information encoded into the genome into morphological form and body plan. However, as our delineation of developmental systems progresses, we are facing increasingly large and complex networks, which cannot be fully and rigorously understood without proper formalisation. This is clearly the case for the GRN governing D/V patterning of the sea urchin embryo, which relies on numerous signalling and regulatory factors, involved in multiples positive and negative feedback circuits.

In our modelling study, several key choices had to be made. As little is known regarding detailed mechanisms and kinetic parameters, we opted for a qualitative, logical formalism. However, to properly model morphogen diffusion and dose-dependent effects, we considered a multilevel extension of the classical Boolean framework. Importantly, in the course of its conception, the model was systematically tested through extensive simulations of wild-type and perturbed conditions. In wild-type conditions, our unicellular model fully recapitulated each territory pattern independently. We further took advantage of a recent multicellular extension of the logical framework to explicitly simulate spatial pattern formation, whose results can be more easily compared directly with the phenotypes of wild-type and morphant embryos.

A key step in our study was to model the interplay between the Nodal and BMP pathways. In this respect, we were guided by our experiments dealing with the treatment of embryos with recombinant Nodal or BMP2/4 proteins at blastula stage (i.e. after the initial specification of the ventral and dorsal territories). These experiments demonstrated that over-activation of one of these pathways is sufficient to abrogate signalling from the other pathway. Consequently, our model highlights the strong antagonism between Nodal and BMP2/4 signalling and suggests that the outcome of this competition relies at least in part on the relative doses of the two TGF-βs, rather than or in addition to their timing of activation. We further suggest that the antagonism stems from a competition for the recruitment of Smad protein complexes. Although evidence for binding site of Smad complexes on the *cis*-regulatory regions of BMP2/4 and Nodal target genes has been described (Hill, 2016), the mechanism of competition between the two cascades has not been fully clarified yet to our knowledge and further experiments are needed to support and validate this hypothesis.

This competition between Nodal and BMP2/4 plays a key role in understanding the regulatory dynamics within the *chordin* knock-down experiments, which was the only morphant not fully recapitulated by our model for all three territories. In the case of the *chordin* knock-down, our logical model predicted that both the ventral and dorsal steady states were possible in the presumptive ventral region. Accordingly, in the *chordin* morphant, both the Nodal and the BMP2/4 pathways are activated, antagonising each other. Following this ectopic activation of BMP signalling, the ventral territory in *chordin* morphants displays a transient dorsalisation, before reversing towards a ventral ectoderm fate during gastrulation, as shown by the presence of a mouth opening. To further explore the underlying regulatory mechanism of this dorsal fate dominance, we performed a stochastic logical simulation of the unicellular model in *chordin* knock-down condition. This analysis resulted in a higher proportion of active dorsal fate over ventral fate, in agreement with the experiments. This result suggests that the transient dorsal dominance is encoded in the structure of the GRN. Indeed, even in the case of the *chordin* morphants, the model accurately recapitulates the conflict caused by the coactivation of the Nodal and BMP2/4 pathways in the ventral ectoderm. However, by modulating the rates associated with the different Smads and performing additional simulation, we showed that the outcome of the competition between the two pathways is sensitive to these rates in the context of co-activation.

At this point, our model remains limited to the major early dorsal-ventral patterning events occurring in sea urchin. However, this model could be tentatively extended in the future to integrate novel data and explore more refined specification and differentiation events. For example, it could be extended to investigate the specification of the boundary ectoderm region, located at the interface between the ectoderm and endomesoderm, which plays a central role in positioning the clusters of primary mesenchyme cells and spicules patterning (Armstrong and McClay, 1994; Armstrong et al., 1993; Duloquin et al., 2007; Hardin et al., 1992; Röttinger et al., 2008). This process is known to depend on Wnt signalling, presumably in conjunction with Nodal, BMP2/4 and ADMP2 signalling (Lapraz et al., 2015; McIntyre et al., 2013; Röttinger et al., 2008; Saudemont et al., 2010). With the current unicellular model, the simulation with the input levels corresponding to the boundary ectoderm (i.e. Admp2 and Wnt active) results in a dorsal stable state (Fig. S2).

Another possible extension to the model would be the integration of the negative feedback of Smad6 on Nodal and BMP2/4 pathways (Armstrong and McClay, 1994; Armstrong et al., 1993; Duloquin et al., 2007; Hardin et al., 1992; Röttinger et al., 2008). Indeed, Smad6 is activated by the dorsal signalling downstream of BMP2/4. Such a negative feedback circuit typically generates an oscillatory behaviour. With the logical formalism, the consideration of this negative circuit would result in a cyclic attractor with alternation of active and inactive BMP and Smad6 activities, which are more difficult to interpret than stable states.

Our logical model focuses on the blastula and gastrula stages of sea urchin embryogenesis. One possible extension would be to further explore the regulatory interactions taking place at earlier stages. In the case of the 32-cell stage, our model correctly recapitulates wild-type pattern mainly driven by Panda expression. Furthermore, the simulation results of Panda loss-of-function in 32-cell stage conditions mirror the fully ventralised phenotype obtained experimentally (Haillot et al., 2015) (Fig. S3 A-B). However, the simulations of Panda overexpression show discrepancies relative to the experimental observations. Indeed, our model predicts a loss of ventral fate specification, whereas global injection of Panda mRNA does not impact the wild-type pattern. Current models suppose that an asymmetry of Panda mRNA provides the spatial cue that in turn controls the polarised activation of downstream genes. Therefore, an asymmetry of *panda* mRNA or of Panda protein constitutes the main driving signal to allocate cell fates, rather than a change in overall Panda concentration. Such signalling based on multicellular gradient cannot be currently recapitulated by our unicellular model, as it requires to integrate inputs from multiple surrounding cells and also to rely on relative differences in concentration instead of absolute levels. As relative concentration differences are tricky to model with the logical formalism, a continuous framework (e.g. ODE) would be better suited to further explore the specificities of such multicellular gradient signalling.

To conclude, using a qualitative, logical model, we could capture several salient dynamical features of the GRN governing the early dorsal-ventral patterning of sea urchin embryos, including the key role played by intercellular interactions. Such qualitative models are useful to explore the interplay between maternal factors and zygotic genes, which orchestrates patterning of the ectoderm of the sea urchin embryo downstream of intercellular signals. To ease further model analyses and extensions, we provide our models as supplementary material, together with a Jupyter notebook implementing all the simulations performed with GINsim and MaBoss (see supplementary material), and a Docker image containing the necessary softwares (https://github.com/colomoto/colomoto-docker).

## Material and methods

### Animals, embryos and treatments

Adult sea urchins (*Paracentrotus lividus*) were collected in the bay of Villefranche-sur-Mer. Embryos were cultured as described in (Lepage and Gache, 1989; Lepage and Gache, 1990). Fertilization envelopes were removed by adding 1mM 3-amino-1,2,4 triazole (ATA) 1 min before insemination to prevent hardening of this envelope followed by filtration through a 75 μm nylon net. Treatments with recombinant BMP2/4 (Recombinant Mouse BMP-4 Protein, CF 5020-BP/CF) or Nodal (Recombinant human Nodal protein 3218-ND-025/CF) proteins were performed at the time indicated in the schemes by adding the recombinant protein diluted from stocks in 1 mM HCl in 24 well plates containing about 1000 embryos in 2 mL of artificial sea water (Saudemont et al., 2010). All the experiments described in this study have been repeated two or three times. At least 200 wild-type and 50 injected embryos were analysed for each condition or experiment. In the case of treatments with recombinant proteins, more than 500 embryos were scored for morphological phenotypes for each condition and about 250 embryos were used for in situ hybridisation. Only phenotypes observed in more than 90% of the embryos are shown.

### Overexpression of mRNAs and morpholino injections

For overexpression studies, capped mRNAs were synthesized from NotI-linearized templates using mMessage mMachine kit (Ambion). After synthesis, capped RNAs were purified on Sephadex G50 columns and quantitated by spectrophotometry. The *nodal*, *lefty*, *chordin* and *bmp2/4* pCS2 constructs have been described in Duboc et al. (2004, 2008) and Lapraz et al. (2009). In this study, *nodal* mRNA was injected at 400 µg/ml, *lefty* mRNA at 200 µg/ml, *chordin* mRNA at 1000 µg/ml and *bmp2/4* mRNA at 500 µg/ml, respectively. Approximately 2-4pL of capped mRNA were injected mixed with Tetramethylrhodamine Dextran (10000 MW) at 5 mg/ml (Duboc et al., 2004). Morpholino oligonucleotides were dissolved in sterile water together with Tetramethylrhodamine Dextran (10000 MW) at 5 mg/ml and approximately 2-4pL of the resulting solution was injected at the one-cell stage. *nodal* morpholino was used at 1mM (Duboc et al., 2004). *chordin* and *bmp2/4* morpholinos were injected at 1mM and 0.2mM, respectively (Lapraz., et al 2009), *lefty* morpholino at 1.5mM (Duboc et al., 2008) and *panda* morpholino at 1.2mM (Haillot et al., 2015).

### Anti-phospho-Smad1/5/8 Immunostaining

Embryos were fixed with 4% formaldehyde for 15 min at swimming blastula stage (3 hours after adding BMP2/4 protein) then briefly permeabilized with methanol. Anti-Phospho- Smad1 (Ser463/465) / Smad5 (Ser463/465) / Smad9 (Ser465/467) from Cell Signalling (D5B10 Ref. 13820) was used at 1/400. Embryos were imaged with an Axio Imager.M2 microscope.

### *In situ* hybridization

*In situ* hybridization was performed using standard methods (Harland, 1991) with DIG-labelled RNA probes and developed with NBT/BCIP reagent. The *nodal*, *chordin* and *tbx2/3* probes have been described previously (Duboc et al., 2004; Lapraz et al., 2009). Control and experimental embryos were developed for the same time in the same experiments. Embryos were imaged with an Axio Imager M2 microscope.

### Logical formalism

We built our model using the multilevel logical formalism introduced by R. Thomas (Thomas and D’Ari, 1990). This qualitative approach relies on graph-based representations of the network and of its dynamics. The network is formalised as a *regulatory graph*, where nodes denote molecular species (e.g. proteins), whereas signed arcs denote regulatory interactions, positive or negative. The nodes can take a limited number of integer values, only two (0 or 1) in the simplest, Boolean case, but potentially more when biologically justified, for example in the case of morphogens with clearly distinct activity ranges. Hence, each regulatory arc is associated with an integer threshold, always 1 in the Boolean case, but potentially higher in the case of a multilevel regulator. Logical rules further specify how each node reacts to the presence or absence of the corresponding incoming interactions. Specific (non-overlapping) Boolean rules are defined for each value of each node.

Boolean rules are built by combining literals (i.e. valued component) with the logic operators AND (denoted “&”), OR (denoted “|”) and NOT (denoted “!”).

Table 1 lists the formula associated with the different components of our model.

Note that the formula associated with zero values are omitted, as they can be computed directly as the complement of the formulae defined for the other values for a given node. For example, the formula of the node FoxA

FoxA => 1 IFF (FoxA | Brachyury) & !Repressor_R1”

can be translated into “FoxA node tends toward the value 1 if and only if FoxA or Brachyury is active and Repressor_R1 is not active”.

In this example, the regulatory actions from Brachyury to FoxA and from Repressor_R1 to FoxA correspond to an activation and to an inhibition, respectively.

The levels of the input (unregulated) nodes are defined at the start of simulations.

Using the Boolean rules of Table 1, we can simulate the behaviour of the system for different input value combinations. In this respect, we use the asynchronous updating approach proposed by R. Thomas (Thomas, 1991), which consists in following all the different possible single unitary changes of values induced by the rules.

The dynamical behaviour of the model is generally represented in terms of a *state transition graph*, where each node represents a discrete state of the model (i.e. a vector listing the values of the different components of the model), whereas each arc represents a state transition.

In this work, we took advantage of the implementation of this logical formalism into the user- friendly Java software suite GINsim (version 3.0, see http://ginsim.org, (Naldi, 2018)). In our analyses, we particularly focused on stable states (see e.g. Fig. 5B), which typically denote cell differentiation states. These can be directly computed (i.e. without unfolding the state transition graph) using a very efficient algorithm implemented in GINsim (Naldi et al., 2007).

### Wild type simulation

We simulated the behaviour of each dorsal-ventral region independently, considering different sets of fixed values for the input nodes in the ventral, ciliary and dorsal presumptive territories. These sets of input values were defined based on previously published results (see Results section). To account for their transient activity, the input nodes for Panda and Admp2 were defined as having a basal value of 0, enabling them to turn off during the simulation if active in the initial state conditions.

As we simulate each territory individually, the unicellular model cannot directly take into account the diffusion of morphogens, which are therefore specified as input levels (e.g. the presence of Lefty is considered as an active input in the ciliary regions, although it is known that it diffuses from the ventral region). For each simulation, we extract the resulting stable state(s) and classify them as ventral, ciliary or dorsal pattern depending on the set of output node levels. For example, the initial conditions for the simulation of the ventral ectoderm territory amount to consider the inputs Nodal (level 2), Lefty, Admp1, Tolloid, Chordin (level 2) and BMP2/4 as active (while the other inputs are inactive). This combination results in a stable state with all the ventral nodes active and the dorsal nodes inactive. In contrast, when the initial conditions are set with Nodal (level 1), Lefty, BMP2/4, Chordin (level 1) and Tolloid being active (and the other inputs inactive), the resulting stable state corresponds to the dorsal fate.

### Morphant simulations

Genetic perturbations are defined in GINsim by constraining the values of selected nodes of the model. To simulate a *knock down* morphant (e.g. injection of a morpholino), the level of the corresponding node is set and maintained to 0. In the case of an ectopic expression (e.g. injection of a mRNA), the level of the corresponding node is set and maintained to its maximal value, which can be 1 or higher in case of a multilevel node. Morphogen diffusion is taken into account through the specifications of proper input values, which thus need to be adjusted for each morphant. For example, the ectopic activation of Nodal is known to induce the activation of its downstream target BMP2/4 very early on; hence, the corresponding input variables must be set at their highest levels for the simulation of ectopic nodal expression.

### Stochastic modelling using MaBoss

When several stable states can be reached (as in the case of *chordin* morpholino), we have performed probabilistic simulations to evaluate the probability to reach each of these stable states from the specified initial conditions. In this respect we used the software MaBoss (https://maboss.curie.fr), a C++ software enabling the simulation of continuous/discrete time Markov processes, applied to Boolean networks. The original unicellular model is converted into the MaBoSS compliant format using a specific export functionality of GINsim, which involves the replacement of multilevel nodes by sets of Boolean variables, without affecting the model dynamic (Stoll et al., 2017). Per default, all up and down rates are considered equal, but these can be modified at will.

In this study, we used MaBoSS to simulate the *chordin* morpholino perturbation (comparing it with the wild-type situation), which resulted in two possible stable states in the unicellular model. The inputs were fixed as for the ventral configuration (Nodal, Lefty, BMP2/4 and Admp1 active) in the presence or inactivation of Chordin. We then modified the propensity to activate the ventral or the dorsal cascade by adjusting the ratios of the rates assigned to the Alk receptors corresponding to each of the two cascades: 0.5/0.5 (equiprobable rates), 0.75/0.25 (ratio favouring the dorsal Alk), 0.25/0.75 (ratio favouring the ventral Alk).

### Multicellular simulation using EpiLog

We took advantage of recent software Epilog (https://epilog-tool.org, v1.1.1.) (Varela et al., 2019) to perform multicellular simulations. Our epithelium model is nine cells wide and six cells long, made of hexagonal shaped cells, each one being in direct contact with at most six different neighbouring cells. The top and bottom part of the epithelium are wrapped together to allow diffusion of signalling molecules through these two sides. Each hexagonal cell encompasses one copy of our unicellular model and thus behave accordingly to the same logical rules during simulations. In contrast with our previous unicellular simulations, the cell inputs are dynamically updated based on the signals perceived from the corresponding output nodes of neighbouring cells (e.g. Nodal input level will be based on the quantity of neighbour cells expressing Nodal as an output). The rules integrating the extracellular signals are identical for all cells of the epithelium. In our epithelium simulations, input nodes of all cells are updated in a synchronous manner. Hence, each epithelium simulation gives rise to a deterministic trajectory ending in a single attractor at the level of the whole tissue (a stable state for the simulations reported here).

The dynamical update of the input node levels implies the definition of additional logical rules for the diffusion of signals, e.g. of the values of input nodes depending on the output nodes active in neighbouring cells, taking into account their distance from the target cell. For example, the rule “{Nodal:2[1:], min = 1}” states that a cell will have its Nodal input node value converging toward the value 1 if at least 1 cell is expressing Nodal at a value 2 at a minimum distance of one cell (i.e. all cell except the target cell itself).

### Multicellular wild type and morphant simulations

For our epithelium simulations, we define the initial state by selecting the nodes that will be active in a specific set of cells at the start of the simulation. During simulations, the values of these nodes can change depending on the model state and on paracrine signalling. To simulate a wild-type embryo, we set the model to an initial state where nodal is broadly expressed at a moderate level (level 1), and where panda is expressed in the presumptive ciliary and dorsal territories (6 rightmost cell columns of the epithelium), with the initial and transient activation of Panda and Nodal (output) nodes. Univin is also ubiquitously present at initial state. As in the unicellular simulation, to simulate loss- or gain-of-function perturbations, the value of the corresponding node is set and maintained at a fixed value. For the siamese simulation, we use the wild-type logical model, with an initial state accounting for a ventral expression of Smad1/4/5/8 and Nodal on the ventral side, but also on the dorsal side of the epithelium (1 rightmost cell column of the epithelium), i.e. a symmetrical activity pattern.

### Model and code availability

The unicellular and multicellular models are available in the GINsim model repository (http://ginsim.org/node/236), together with the Jupyter notebook encoding all the simulations performed with GINsim and MaBoss, which is also available in a GitHub repository (https://github.com/ComputationalSystemsBiology/echinodal_notebook). The Jupyter notebook uses the colomoto-docker image (https://github.com/colomoto/colomoto-docker, v2020-01-24) (Naldi et al., 2018). The models can be uploaded in zginml and peps format, to be open with GINsim (v3.0.0b) and EpiLog (v1.1.1), respectively. The unicellular model has been further deposited in SBML qual format in the BioModels database (ID MODEL2002190001), together with its reference annotations.

## Acknowledgements

We thank Aline Chessel for excellent technical help. We are indebted to Guillaume Lavisse for insightful comments. We thank Aurélie Martres for taking care of the sea urchins. We thank Aurelien Naldi for continuous help and support on the development of the Colomoto Jupyter Notebook. We thank Claudine Chaouiya and Pedro Monteiro for their support regarding the use of EpiLog. We thank Mathurin Dorel for his help in defining a preliminary version of the cellular model.

## Funding

This work was supported by grants from the CNRS, the UNSA, the French Foundation for Cancer Research (ARC) SFI20121205586, and the National Agency for Research (ANR) (grant Echinodal ANR-14-CE11-0006-01 to TL), and by the FRM Team project DEQ20180339195 to TL. S. F. has been supported by a grant from the PhD School *Life Science Complexity* (ED 515) and another grant from the *Fondation pour la Recherche Médicale* FDT201904008366. M.D. Molina was supported by an EMBO long-term fellowship and by an ARC postdoctoral fellowship. E. Haillot was supported by a 4th year doctoral fellowship from the ARC. M.T.C. has been further supported by the *Institut Universitaire de France*.

## Competing interests

No competing interests declared.

---

**Supplementary Figure 1.**
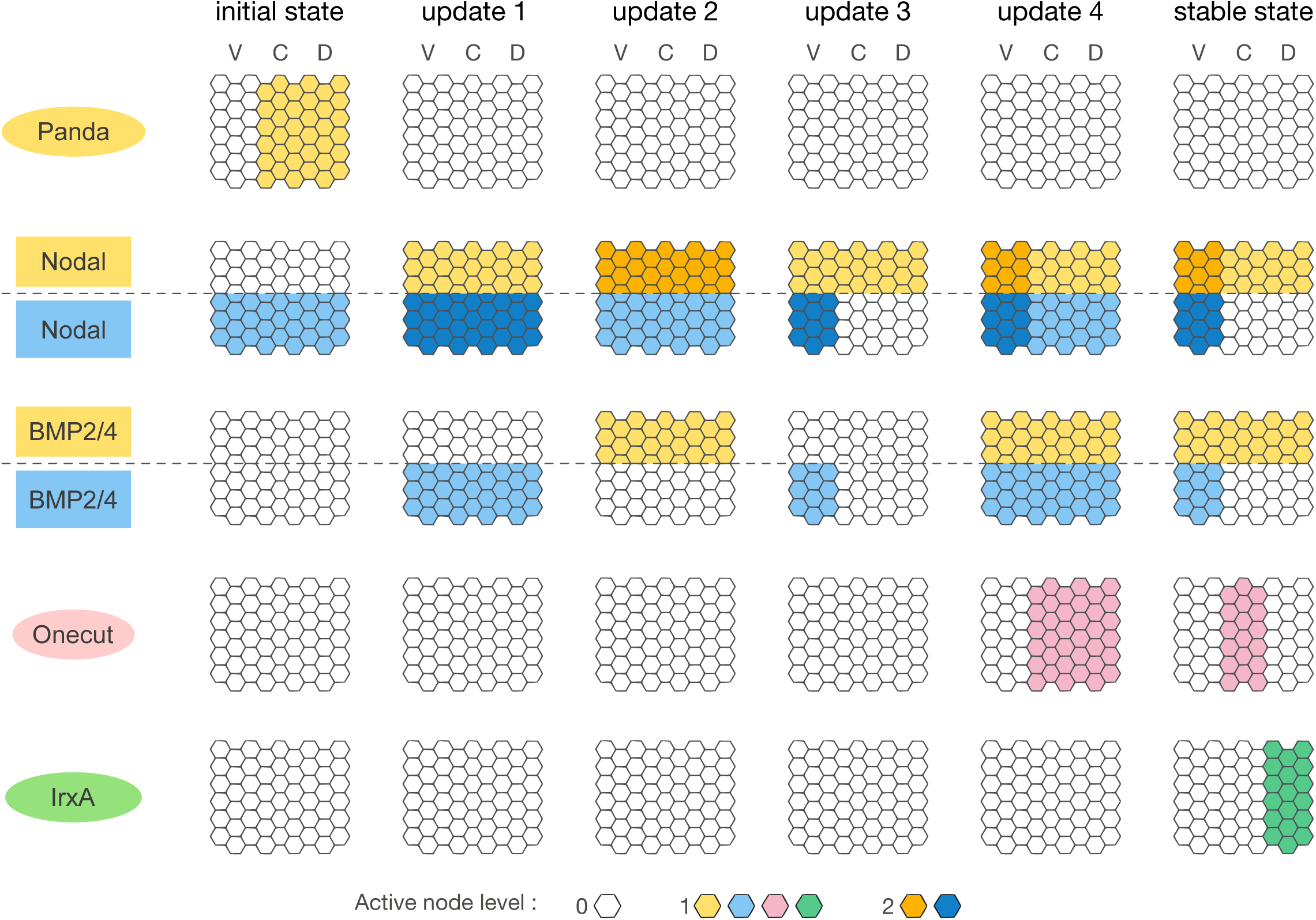
Intermediary states reached during wild-type simulation with EpiLog. Starting from the initial state (left) with Nodal ubiquitously active and Panda restricted to the ciliary and dorsal presumptive territories, the Epilog simulation of the wild-type condition first predicts a transient expression of *nodal* in the whole ectoderm (Nodal output node in dark and light blue), which is then restricted to the ventral region in the stable state (right), although Nodal proteins still diffuse toward the dorsal region (Nodal input node in yellow). In parallel, the *bmp2/4* expression becomes restricted to the ventral side (BMP2/4 output node in blue) and BMP2/4 protein diffuse toward the dorsal region (BMP2/4 input node in yellow). The activity dynamics of the two markers genes *onecut* (pink, ciliary) and *irxa* (green, dorsal) denotes the progressive restriction of the ciliary band in the central region, as both Nodal and BMP2/4 cascade takes place in the ventral and dorsal territories. For the multilevel node Nodal, higher activity level is depicted by darker colours.

**Supplementary Figure 2.**
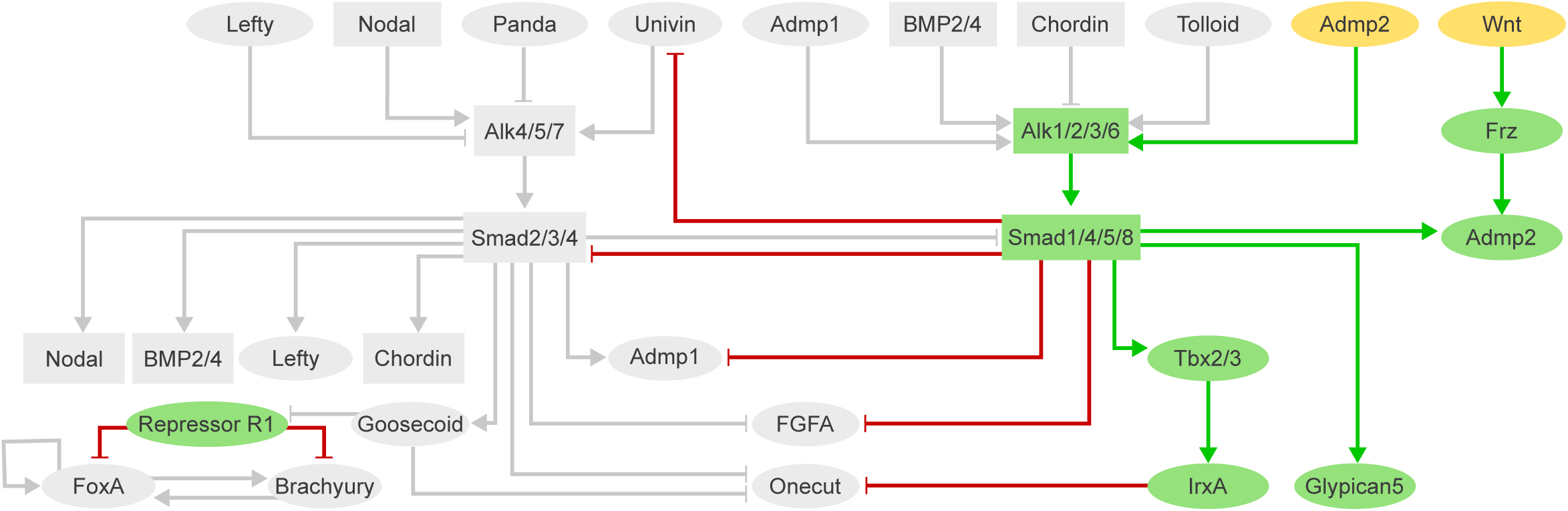
Simulation of the boundary ectoderm in the unicellular model. Stable state obtained with the unicellular model when considering Admp2 and Wnt inputs active. Active nodes (yellow for inputs and green for dorsal nodes) and edges (green for activation and red for inhibition) are shown in colour, inactive ones are shown in grey. This stable state mirror corresponds exactly to the dorsal stable state shown in Figure 6.

**Supplementary Figure 3.**
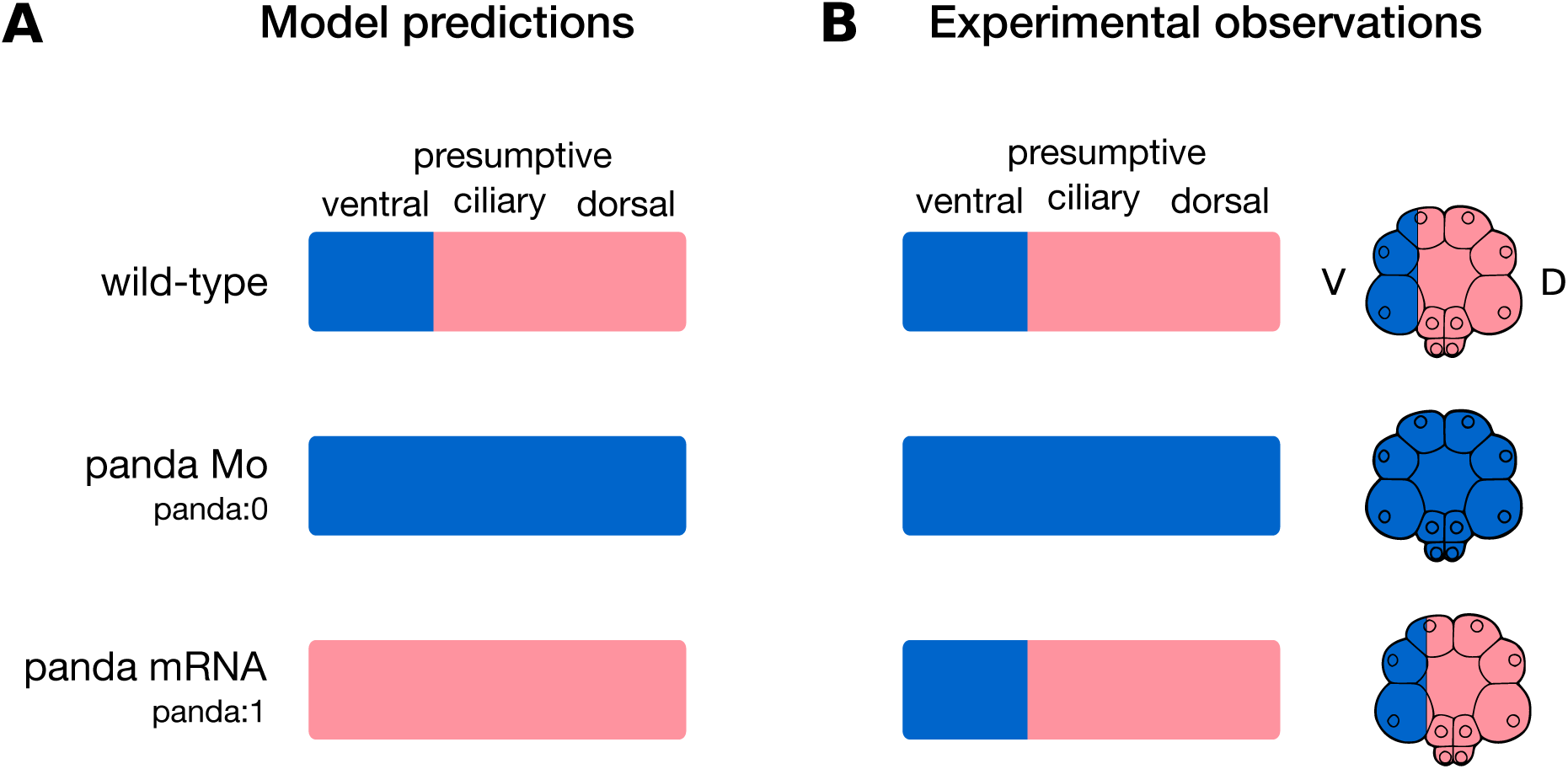
Simulation of *panda* perturbations at the 32-cell stage. Starting with a restricted combination of inputs with Nodal and Panda, we simulated the 32- cell stage D/V patterning using the unicellular model. In wild-type condition, model predictions (A) correctly recapitulates experimental observations (B). It is also the case for the simulation of Panda loss of function, mirroring the fully ventralised phenotype observed upon *panda* Mo injection. Simulations of *panda* overexpression fully abrogate ventral specification and result in a global ciliary phenotype (A), whereas experimental evidences show no impact of global overexpression of *panda* on the onset of D/V patterning (Haillot et al., 2015) (B).

## Notes

http://ginsim.org/node/236

### Competing Interest Statement

The authors have declared no competing interest.

### Summary of Updates

The model has been improved and the manuscript updated accordingly.

## References

1. Armstrong, N. and McClay, D. R. (1994). Skeletal pattern is specified autonomously by the primary mesenchyme cells in sea urchin embryos. Dev. Biol. 162, 329–338.

2. Armstrong, N., Hardin, J. and McClay, D. R. (1993). Cell-cell interactions regulate skeleton formation in the sea urchin embryo. Dev. Camb. Engl. 119, 833–840.

3. Arnone, M. I. and Davidson, E. H. (1997). The hardwiring of development: organization and function of genomic regulatory systems. Development 124, 1851–1864.

4. Ben-Zvi, D., Shilo, B.-Z., Fainsod, A. and Barkai, N. (2008). Scaling of the BMP activation gradient in Xenopus embryos. Nature 453, 1205–1211.

5. Bolouri, H. and Davidson, E. H. (2010). The gene regulatory network basis of the “community effect,” and analysis of a sea urchin embryo example. Developmental Biology 340, 170–178.

6. Chaouiya, C., Naldi, A. and Thieffry, D. (2012). Logical modelling of gene regulatory networks with GINsim. Methods Mol. Biol. Clifton NJ 804, 463–479.

7. Chen, D., Zhao, M. and Mundy, G. R. (2004). Bone Morphogenetic Proteins. Growth Factors 22, 233–241.

8. Cheng, S. K., Olale, F., Brivanlou, A. H. and Schier, A. F. (2004). Lefty Blocks a Subset of TGFβ Signals by Antagonizing EGF-CFC Coreceptors. PLOS Biol. 2, e30.

9. De Robertis, E. M. (2009). Spemann’s organizer and the self-regulation of embryonic fields. Mech. Dev. 126, 925–941.

10. Duboc, V., Röttinger, E., Besnardeau, L. and Lepage, T. (2004). Nodal and BMP2/4 signaling organizes the oral-aboral axis of the sea urchin embryo. Dev. Cell 6, 397–410.

11. Duboc, V., Lapraz, F., Besnardeau, L. and Lepage, T. (2008). Lefty acts as an essential modulator of Nodal activity during sea urchin oral-aboral axis formation. Dev. Biol. 320, 49–59.

12. Duloquin, L., Lhomond, G. and Gache, C. (2007). Localized VEGF signaling from ectoderm to mesenchyme cells controls morphogenesis of the sea urchin embryo skeleton. Dev. Camb. Engl. 134, 2293–2302.

13. Haillot, E., Molina, M. D., Lapraz, F. and Lepage, T. (2015). The Maternal Maverick/GDF15-like TGF-β Ligand Panda Directs Dorsal-Ventral Axis Formation by Restricting Nodal Expression in the Sea Urchin Embryo. PLoS Biol. 13, 1–38.

14. Hardin, J., Coffman, J. A., Black, S. D. and McClay, D. R. (1992). Commitment along the dorsoventral axis of the sea urchin embryo is altered in response to NiCl2. Dev. Camb. Engl. 116, 671–685.

15. Harland, R. M. (1991). In situ hybridization: an improved whole-mount method for Xenopus embryos. Methods Cell Biol. 36, 685–695.

16. Hill, C. S. (2016). Transcriptional Control by the SMADs. Cold Spring Harb Perspect Biol 8,.

17. Joubin, K. and Stern, C. D. (2001). Formation and maintenance of the organizer among the vertebrates. Int. J. Dev. Biol. 45, 165–175.

18. Juan, H. and Hamada, H. (2001). Roles of nodal-lefty regulatory loops in embryonic patterning of vertebrates. Genes Cells Devoted Mol. Cell. Mech. 6, 923–930.

19. Kimelman, D. and Pyati, U. J. (2005). Bmp signaling: turning a half into a whole. Cell 123, 982–984.

20. Lapraz, F., Röttinger, E., Duboc, V., Range, R., Duloquin, L., Walton, K., Wu, S.-Y., Bradham, C., Loza, M. A., Hibino, T., et al. (2006). RTK and TGF-beta signaling pathways genes in the sea urchin genome. Dev. Biol. 300, 132–152.

21. Lapraz, F., Besnardeau, L. and Lepage, T. (2009). Patterning of the dorsal-ventral axis in echinoderms: Insights into the evolution of the BMP-chordin signaling network. PLoS Biol. 7,.

22. Lapraz, F., Haillot, E. and Lepage, T. (2015). A deuterostome origin of the Spemann organiser suggested by Nodal and ADMPs functions in Echinoderms. Nat. Commun. 6, 8434.

23. Lele, Z., Nowak, M. and Hammerschmidt, M. (2001). Zebrafish admp is required to restrict the size of the organizer and to promote posterior and ventral development. Dev. Dyn. 222, 681–687.

24. Lepage, T. and Gache, C. (1989). Purification and characterization of the sea urchin embryo hatching enzyme. J. Biol. Chem. 264, 4787–4793.

25. Lepage, T. and Gache, C. (1990). Early expression of a collagenase-like hatching enzyme gene in the sea urchin embryo. EMBO J. 9, 3003–3012.

26. Levine, M. and Davidson, E. H. (2005). Gene regulatory networks for development. Proc. Natl. Acad. Sci. 102, 4936–4942.

27. Li, E., Materna, S. C. and Davidson, E. H. (2012). Direct and indirect control of oral ectoderm regulatory gene expression by Nodal signaling in the sea urchin embryo. Dev. Biol. 369, 377–385.

28. Li, E., Materna, S. C. and Davidson, E. H. (2013). New regulatory circuit controlling spatial and temporal gene expression in the sea urchin embryo oral ectoderm GRN. Dev. Biol. 382, 268–279.

29. Li, E., Cui, M., Peter, I. S. and Davidson, E. H. (2014). Encoding regulatory state boundaries in the pregastrular oral ectoderm of the sea urchin embryo. Proc. Natl. Acad. Sci. 111, E906–E913.

30. Mbodj, A., Gustafson, E. H., Ciglar, L., Junion, G., Gonzalez, A., Girardot, C., Perrin, L., Furlong, E. E. M. and Thieffry, D. (2016). Qualitative Dynamical Modelling Can Formally Explain Mesoderm Specification and Predict Novel Developmental Phenotypes. PLoS Comput. Biol. 12, 1–17.

31. McIntyre, D. C., Seay, N. W., Croce, J. C. and McClay, D. R. (2013). Short-range Wnt5 signaling initiates specification of sea urchin posterior ectoderm. Dev. Camb. Engl. 140, 4881–4889.

32. Molina, M. D. and Lepage, T. (2020). Maternal factors regulating symmetry breaking and dorsal–ventral axis formation in the sea urchin embryo. In Current Topics in Developmental Biology, pp. 283–316. Elsevier.

33. Molina, M. D., Quirin, M., Haillot, E., De Crozé, N., Range, R., Rouel, M., Jimenez, F., Amrouche, R., Chessel, A. and Lepage, T. (2018). MAPK and GSK3/ß-TRCP- mediated degradation of the maternal Ets domain transcriptional repressor Yan/Tel controls the spatial expression of nodal in the sea urchin embryo. PLoS Genet. 14, e1007621.

34. Molina, M. D., de Crozé, N., Haillot, E. and Lepage, T. (2013). Nodal: master and commander of the dorsal–ventral and left–right axes in the sea urchin embryo. Current Opinion in Genetics & Development 23, 445–453.

35. Naldi, A. (2018). BioLQM: A Java Toolkit for the Manipulation and Conversion of Logical Qualitative Models of Biological Networks. Front. Physiol. 9, 1605.

36. Naldi, A., Thieffry, D. and Chaouiya, C. (2007). Decision Diagrams for the Representation and Analysis of Logical Models of Genetic Networks. In Computational Methods in Systems Biology (ed. Calder, M.) and Gilmore, S.), pp. 233–247. Berlin, Heidelberg: Springer.

37. Peter, I. S., Faure, E. and Davidson, E. H. (2012). Predictive computation of genomic logic processing functions in embryonic development. Proc. Natl. Acad. Sci. 109, 16434– 16442.

38. Range, R. and Lepage, T. (2011). Maternal Oct1/2 is required for Nodal and Vg1/Univin expression during dorsal–ventral axis specification in the sea urchin embryo. Developmental Biology 357, 440–449.

39. Range, R., Lapraz, F., Quirin, M., Marro, S., Besnardeau, L. and Lepage, T. (2007). Cis-regulatory analysis of nodal and maternal control of dorsal-ventral axis formation by Univin, a TGF-β related to Vg1. Development 134, 3649–3664.

40. Reversade, B. and De Robertis, E. M. (2005). Regulation of ADMP and BMP2/4/7 at opposite embryonic poles generates a self-regulating morphogenetic field. Cell 123, 1147–1160.

41. Röttinger, E., Saudemont, A., Duboc, V., Besnardeau, L., McClay, D. and Lepage, T. (2008). FGF signals guide migration of mesenchymal cells, control skeletal morphogenesis [corrected] and regulate gastrulation during sea urchin development. Dev. Camb. Engl. 135, 353–365.

42. Sakuma, R., Ohnishi, Y., Meno, C., Fujii, H., Juan, H., Takeuchi, J., Ogura, T., Li, E., Miyazono, K. and Hamada, H. (2002). Inhibition of Nodal signalling by Lefty mediated through interaction with common receptors and efficient diffusion. Genes Cells 7, 401–412.

43. Saudemont, A., Haillot, E., Mekpoh, F., Bessodes, N., Quirin, M., Lapraz, F., Duboc, V., Röttinger, E., Range, R., Oisel, A., et al. (2010). Ancestral Regulatory Circuits Governing Ectoderm Patterning Downstream of Nodal and BMP2/4 Revealed by Gene Regulatory Network Analysis in an Echinoderm. PLOS Genet. 6, e1001259.

44. Stoll, G., Caron, B., Viara, E., Dugourd, A., Zinovyev, A., Naldi, A., Kroemer, G., Barillot, E. and Calzone, L. (2017). MaBoSS 2.0: an environment for stochastic Boolean modeling. Bioinformatics 33, 2226–2228.

45. Su, Y.-H., Li, E., Geiss, G. K., Longabaugh, W. J. R., Krämer, A. and Davidson, E. H. (2009). A perturbation model of the gene regulatory network for oral and aboral ectoderm specification in the sea urchin embryo. Dev. Biol. 329, 410–421.

46. Thomas, R. (1991). Regulatory networks seen as asynchronous automata: A logical description. J. Theor. Biol. 153, 1–23.

47. Thomas, R. and D’Ari, R. (1990). Biological feedback. Boca Raton, FL etc.: CRC Press.

48. Varela, P. L., Ramos, C. V., Monteiro, P. T. and Chaouiya, C. (2019). EpiLog: A software for the logical modelling of epithelial dynamics. F1000Research 7, 1145.

49. Wilczynski, B. and Furlong, E. E. M. (2010). Challenges for modeling global gene regulatory networks during development: Insights from Drosophila. Dev. Biol. 340, 161–169.

50. Willot, V., Mathieu, J., Lu, Y., Schmid, B., Sidi, S., Yan, Y.-L., Postlethwait, J. H., Mullins, M., Rosa, F. and Peyriéras, N. (2002). Cooperative Action of ADMP- and BMP-Mediated Pathways in Regulating Cell Fates in the Zebrafish Gastrula. Dev. Biol. 241, 59–78.

